# Lateral interactions override nucleotide state in determining FtsZ filament curvature

**DOI:** 10.64898/2026.07.05.736574

**Authors:** Soumyajit Dutta, Joyeeta Chakraborty, Ekaterina G. Kholina, Ilya B. Kovalenko, Nikita B. Gudimchuk, Pananghat Gayathri

## Abstract

The bacterial tubulin homolog FtsZ assembles into dynamic filaments that form the cytokinetic Z-ring and drives constriction during cell division. Whether a nucleotide-dependent FtsZ filament curvature plays a role in constriction is often debated. Here, we combine cryo–electron microscopy and molecular dynamics simulations to understand the structural basis of FtsZ filament curvature. Cryo-EM structures of GTP-bound Spiroplasma FtsZ filaments in two curved states emphasize that curvature is an intrinsic property of FtsZ filament, confirming recent models in which GTP hydrolysis does not dictate protofilament bending in the tubulin family. Consistently, molecular dynamics simulations demonstrate that GTP-bound filaments can adopt a range of curved conformations. The preferred intrinsic curvature appears to be such that the C-terminal end of the globular domain faces the convex surface. Structural analyses of the curved conformations identify dynamic and stationary zones at the longitudinal interfaces of the protofilament, suggesting that structural plasticity of the intermonomer interface contributes to filament bending. Furthermore, we demonstrate that lateral interactions between adjacent protofilaments straighten the filaments, overriding their relaxed curved states. Optimal orientations of lateral interactions in the Z-ring assembly could be brought about by other interacting proteins of the divisome machinery. The straighter filament conformation is likely to stimulate a higher GTPase activity. Together, our findings establish lateral association as a primary determinant for straight FtsZ filaments, analogous to the tubulin protofilaments in a microtubule lattice. The snapshots of structural states provide a mechanistic basis for how the intrinsic curvature facilitates association on the membrane and the physiological relevance of transitions between bent and straight conformations of the FtsZ filament during Z-ring assembly and constriction.

## Introduction

FtsZ is an essential and highly conserved cell division protein in bacteria. It is a tubulin homolog which polymerizes in the presence of GTP. However, unlike microtubules, FtsZ exhibits treadmilling within the cells, which is dependent on the disassembly of monomers coupled with GTP hydrolysis. Since the discovery of FtsZ, its role in force generation during cell division remains one of the major questions in the field. Coltharp et al. 2016 showed that the rate of septum closure in a dividing *E. coli* cell depends on cell wall synthesis activity rather than the GTPase activity of FtsZ. Additionally, the maximum force generated by the Z-ring was estimated to be 8-80 pN range which is not enough to overcome turgor pressure of 50-300 nN (Whatmore and Reed, 1990, Lan et al., 2009, Deng et al., 2011, Erickson et al., 2010, Hsin et al., 2012, Xiao and Goley, 2016). Thus, how the Z-ring can derive sufficient force for cell constriction remain elusive.

There are two major models proposed for the force generation mechanism: i) filament bending model, and ii) filament sliding model (Szwedziak and Ghosal, 2017). In the bending model, FtsZ in the presence of GTP polymerizes into straight filaments which then hydrolyze GTP to bend and consequently deform the membrane. In the sliding model, short FtsZ filaments overlap via lateral interaction to form a complete ring which then tightens by maximising overlap, leading to membrane constriction. Simulation-based studies have been performed to quantitatively assess these mechanisms. One of the recent studies by Nguyen et al., 2021 determined the necessary conditions for each model to generate constriction force. Because of the lack of a unified force generation mechanism by FtsZ, a few open questions that currently exist are: (i) is there a correlation between filament curvature and GTP-hydrolysis? (ii) what is a preferred intrinsic curvature, if any, of filament bending? (iii) how does treadmilling and protofilament bending dynamically couple to generate force?

The idea of a mechanical force generated by FtsZ filament came from earlier negative staining transmission electron microscopy observations by Lu et al., 2000 suggesting that GTP-bound filaments favored a straight conformation while GDP-bound was curved. Later, *in vitro* reconstitution experiments with *E. coli* FtsZ showed membrane constriction in liposomes when FtsZ was modified with a membrane binding motif, suggesting force generation during filament assembly (Osawa et al., 2008). Importantly, the curvature (concave or convex) of membrane deformation was dependent on the position of the membrane binding motif with respect to the FtsZ filament indicating an intrinsic curvature of the filament (Osawa et al., 2009).

Molecular dynamics simulation studies also proposed the idea of GDP bound filament being more bent compared to GTP filament (Hsin et al., 2012). An analogous concept was also prevalent in the microtubule field where the disassembling microtubule end has a ‘ram’s horn like structure’ due to the curvature of the GDP bound protofilament (Mandelkow et al., 1991). However, more recent studies show that GTP-bound tubulin addition also goes through a step of curved GTP tubulin dimers (Brouhard and Rice, 2018, Gudimchuk and McIntosh, 2021).

Recent investigations have proposed that treadmilling of FtsZ filament can generate torsional stress upon GTP hydrolysis which can deform membrane and this is essential for cell constriction (Ramirez-Diaz et al., 2021, Whitley et al., 2021). Recent models support the idea that during treadmilling the Z-ring generates an inward force that is essential for correct positioning of the peptidoglycan synthesis enzymes at the septum (Whitley et al., 2021, Yang and Liu, 2022). This inward membrane deformation is possibly essential to change the direction of the new PG synthases at the septum (Nguyen et al., 2021). According to the current understanding, treadmilling dynamics of FtsZ filaments helps encounter each other laterally, giving rise to a condensed narrow ring. Then the condensed Z-ring acts as a scaffold to recruit PG synthesis enzymes and guide them to construct an evenly distributed new septum (Walker et. al., 2020, Squyres et al., 2021, Whitley et al., 2021). FtsZ treadmilling has also been found to be required until the initiation of constriction and to become dispensable in the later stages. However, treadmilling has been observed to increase the rate of constriction (Bisson-Filho et al., 2017). Understanding the FtsZ filament dynamics of a cell wall less bacterium such as Spiroplasma will help understand the role of treadmilling in the absence of peptidoglycan synthesis, prompting us to initiate studies on Spiroplasma FtsZ (Chakraborty, et al., 2024) and to explore unique features of Mollicute FtsZs (Dutta, et al., 2025).

Though recent advancement of high-resolution microscopy experiments and simulation studies have greatly improved our understanding of the Z-ring dynamics in vivo, we still do not have a consensus model of force generation mechanism by FtsZ filament. Moreover, the role of membrane anchoring proteins like FtsZ, ZipA, beyond being a passive anchor to the membrane, is not well understood. Earlier experiments on liposomes suggested that full constriction of vesicles could only be achieved in the presence of FtsA* (mutant of FtsA that bypasses the requirement of ZipA) (Osawa and Erickson, 2013). FtsA might play both a stabilizing and anchoring role for FtsZ filaments to effectively transduce the mechanical energy from GTP hydrolysis and relay to the membrane.

One of the promising methods for obtaining high resolution native structure of protein filaments is cryo electron microscopy. However, because of the flexible nature of the FtsZ single protofilament, obtaining atomic resolution structures has been challenging. Hence, high resolution information about the filament conformation of FtsZ has been from crystal structures of FtsZ, in which the crystal packing adopts a filament-like packing (Ruiz et al., 2023). One of the first attempts of capturing *Escherichia coli* FtsZ (with GMPCPP, a slowly hydrolysable GTP analog) in filament state by cryo-EM was limited to a resolution of 7-8 Å. Similarly, the cryo-EM structure of *Mycobacterium tuberculosis* FtsZ (with GMPCPP) filament has been determined at about 7 Å resolution (Wagstaff 2023). The primary reasons for low resolution could be filament flexibility and low twist of the helical filament accompanied by the preferred orientation problem of the filament in vitrified ice (Wagstaff et al. 2017, Wagstaff et al. 2023). Recently, a high-resolution filament structure of FtsZ from *Klebsiella pneumoniae* (KpFtsZ) bound to GMPPCP, a non-hydrolysable analog, has been captured at an overall resolution of 3 Å (Fujita et al., 2023) and of FtsZ1 from Odinarcheaota to 3.6 Å (Kumari, Uthaman et al., 2025), both of which are straight filament conformations with negligible helical twist. Another cryo-EM structure of the FtsZ double protofilament (GMPCPP bound) in complex with ZapA has been reported at 2.73 Å resolution (Fujita et al., 2025). This double protofilament arrangement of FtsZ-ZapA structure primarily revealed details of lateral interactions of FtsZ protofilaments and the effect of ZapA binding in protofilament assembly.

Here, we captured multiple curved conformations of *Spiroplasma melliferum* FtsZ (SmFtsZ) filament bound to GTP, using cryo-electron microscopy without imposing helical symmetry, at a resolution of 3.4 Å and 3.9 Å for two different curvature states respectively. We provide complementary molecular dynamics simulations which correlate with the filament conformations observed using cryo-EM. Our results support an intrinsic curvature for the FtsZ protofilaments in the presence of GTP. The exposure of the C-terminal end of the globular domain to the convex exterior of the curved state demonstrates that the intrinsic curvature is compatible with membrane association along the diameter of the cell. Lateral interaction, either through bundling of filaments at high concentrations in an in vitro experiment or the presence of interacting partners such as FtsA or ZapA within the cell, leads to straightening of the filaments. The bent to straight transitions accompanying the interplay between ring assembly and treadmilling could contribute to the force generation during ring constriction.

## Results

### FtsZ protofilaments intrinsically adopt a curved conformation

To determine whether FtsZ filament curvature is correlated to GTP hydrolysis, we first analysed transmission electron micrographs of negatively stained FtsZ filaments. We measured the curvature of wild type-SmFtsZ and compared it with its hydrolysis deficient mutant-SmFtsZ^D211G^ in the presence of GTP. SmFtsZ^D211G^ filaments formed curved single protofilaments similar to the wild type (Fig. 1A), indicating that GTP hydrolysis is not the primary determinant for the observed curvature. Filaments formed by SmFtsZ in presence of GTP also showed a comparable degree of curvature with values of 25.29 ± 7.72 µm^-1^ for SmFtsZ (mean ± SD, N=70) and 24.65 ± 5.17 µm^-1^ for SmFtsZD^211G^ (mean ± SD, N=56) (Fig. 1B). Thus, the presence of γ-phosphate is not sufficient to enforce a straight protofilament conformation. To check if the filament curvature is not an artifact of sample preparation in negative stain TEM, we estimated the curvature of SmFtsZ filaments from cryo-EM micrographs and confirmed that similar curvature (29.28 ± 9.71 µm^-1^, mean ± SD, N= 69) exists in native vitrified conditions too (Fig S1 A, B).

**Figure 1.**
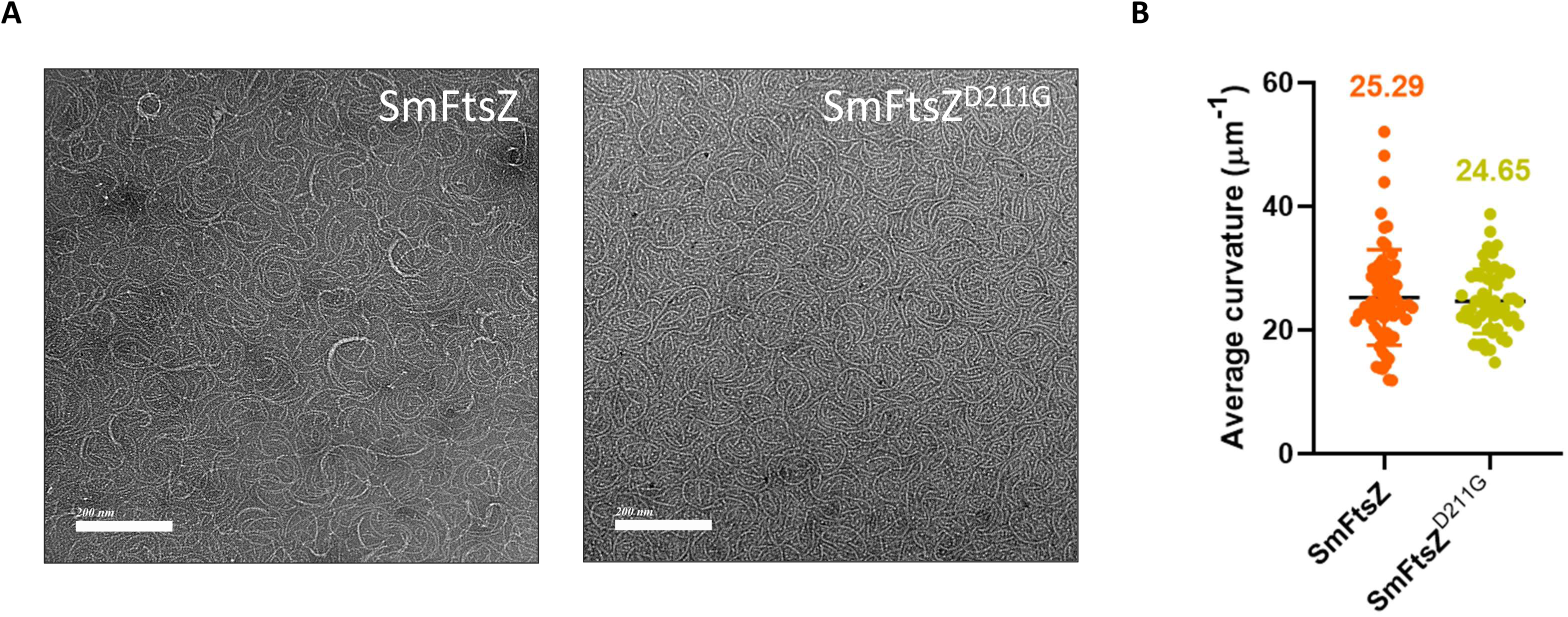
FtsZ protofilament curvature is independent of GTP hydrolysis. A. Negative-stain TEM micrographs of SmFtsZ and SmFtsZ^D211G^ filaments assembled in the presence of GTP. Individual protofilaments display pronounced curvature in both the samples. B. Quantification of filament curvature showing average curvature and mean of the measured filaments. The curvature was calculated for individual filaments as described in the Methods. The bar represents the mean ± SD, and the number above denotes the mean value.

Together our curvature analysis data from FtsZ filaments formed under hydrolysis deficient conditions demonstrate that GTP-bound filaments are curved, rather than straight.

### Cryo-EM reconstruction reveals curved states in SmFtsZ filament with bound GTP

To elucidate the structural basis of curvature in a hydrolysis competent state, we employed single particle cryo-EM to analyze SmFtsZ filaments architecture assembled in the presence of GTP. We obtained high resolution 3D reconstructions of two distinct filament conformations which do not satisfy a helical symmetry or a straight conformation (upon heterogenous refinement of all the particles in same data set), hereafter referred to as curved state 1 (curvature of ∼3° or 11.6 µm^-1^) and state 2 (curvature of ∼7° or 27 µm^-1^) (Fig. 2A, B). The detailed cryo-EM data processing workflow and model refinement statistics have been described in Fig. S2 and Supplementary table 1. The resolution achieved for both the reconstructions was sufficient to resolve secondary structural features unambiguously for 4 monomers of FtsZ filament (state 1 at 3.4 Å and state 2 at 4 Å, FSC threshold 0.143). Density corresponding to the γ-phosphate enabled us to assign GTP in the active site for both state 1 and state 2 reconstructions (Fig. S3; local resolution in the range of ∼2.8 Å and 3.5 Å for state 1 and state 2 respectively). It is important to note that the particles contributing to the two reconstructed maps are heterogeneous in conformation as revealed by 3D variability analysis which possibly limit achieving higher resolution maps (Supplementary movie 1, 2).

**Figure 2.**
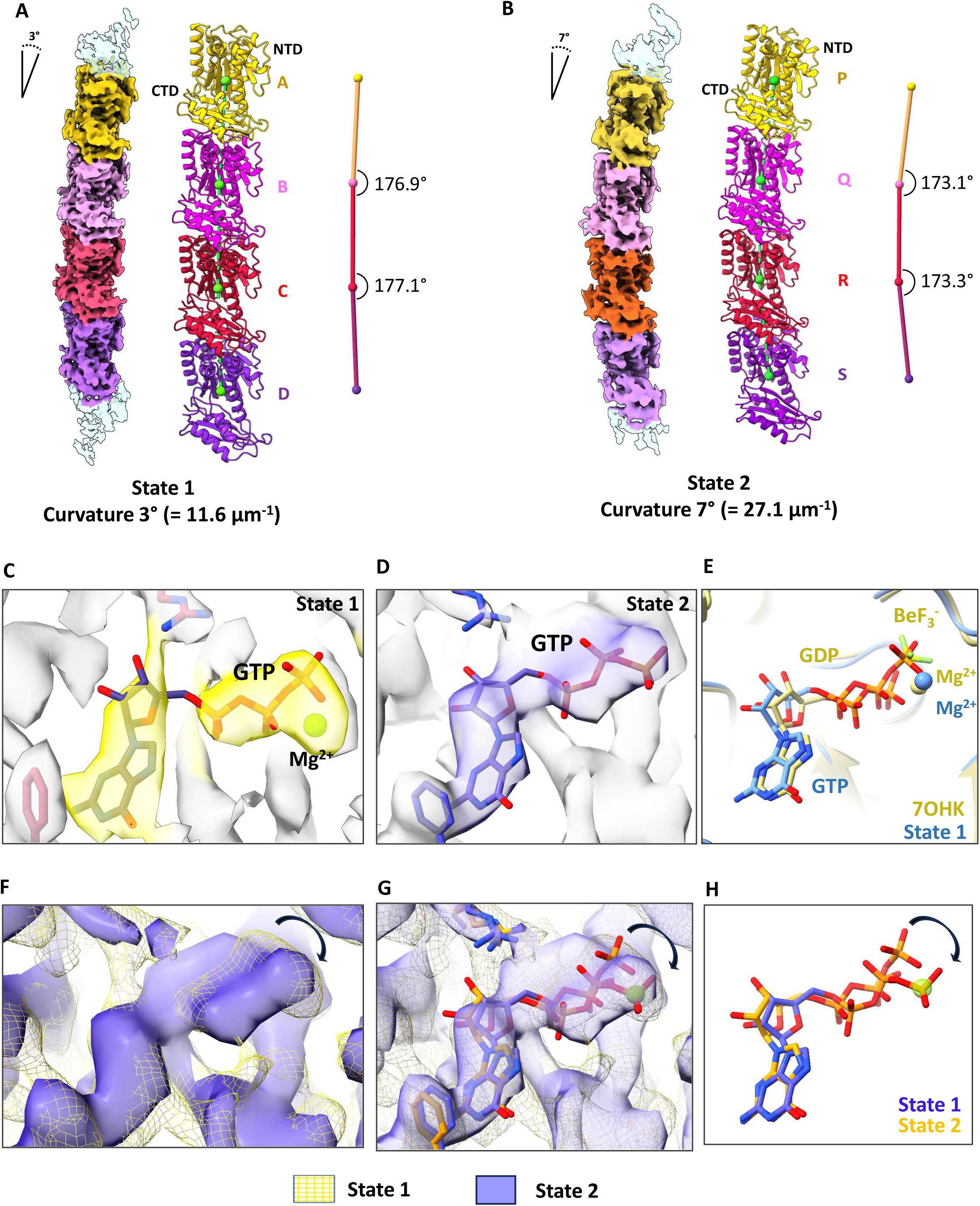
Cryo-EM structure of SmFtsZ in two curvature states. A. 3D map (left), atomic model (middle) and curvature measurements (right) is shown for curved state 1. Centroids of monomers are depicted as spheres connected by straight lines, and the angle between the lines are labelled. B. 3D map (left), atomic model (middle) and curvature measurements (right) is shown for curved state 2. NTD and CTD are marked for the atomic models. C, D. GTP conformation and the corresponding maps for state 1 (C) and state 2 (D). (E) Comparison of the GTP and Mg^2+^ orientation between state 1 structure (blue) with high resolution crystal structure PDB ID 7OHK. (F) Superimposition of the GTP density between state 1 (yellow mesh) and state 2 (purple volume). The movement in the gamma phosphate position is highlighted with an arrowhead. (G, H) Superimposition of the GTP between state 1 (purple) and state 2 (yellow) models.

Importantly, the two curvature states showed differences in conformation of the bound GTP (Fig. 2C-H). The density for GTP in the curved state 1 structure resembles the conformation observed in FtsZ crystal structure from *Staphylococcus aureus* (PDB 7OHK) or *Klebsiella pneumoniae* double FtsZ-ZapA double filament complex solved by cryo-EM (PDB 9ISK) (Fig. 2, S4A), and hence a GTP and a Mg^2+^ ion was modelled in the density. The resolution, and hence the ambiguous density, precludes modelling of the water molecules that coordinate the Mg^2+^ ion. However, for the curved state 2 model, we observe an apparent shift of the GTP γ-phosphate toward the canonical Mg²□ binding site. This conformation is similar to the GMPCPP conformation in the cryo EM structure of Odin FtsZ1 (PDB 9V7V; Kumari and Uthaman et al. 2025) (Fig. S4B). A conformation of GDP and phosphate modelled in the microtubule structure (PDB 6EVX) also roughly matches the position of the γ-phosphate in the curved state 2. However, in our structural states, the γ-phosphate was clearly not detached from the β-phosphate based on distance criteria. The contour levels for density for the three phosphates were also comparable, indicating that the γ-phosphate has not been released in this state. These observations raise the possibility that the γ-phosphate may sample multiple conformations and that filament curvature could influence its positional flexibility or vice versa.

We observed that the direction of curvature in FtsZ filament is such that the N-terminal domain is present towards the concave side (Fig 2A, B), directing the C-terminal end of the globular domain towards the convex surface. This is consistent with the proposal from in vitro studies by Osawa et al. 2009. It is interesting to note that the direction of curvature is opposite to the protofilament bending direction from a disassembling microtubule at the minus end, which is toward the outer side of the lumen. For comparison, we superimposed SmFtsZ curved state 2 with microtubule 13 protofilament structure (PDB 6O2S) and showed that the direction of FtsZ is towards the lumen side with a tilt towards NTD (Fig. S5, Supplementary movie 3).

### Bending of FtsZ filament is driven by dynamic interactions at the longitudinal interface

To characterize the difference in relative orientations of NTD and CTD domains between the two curved states, we superimposed NTDs (or CTDs) of the two models and calculated RMSDs of C^α^ atoms for the CTDs (or the NTDs when CTDs are superimposed) (Fig. S6A). Interestingly, the curvature arises without large-scale rearrangements within the FtsZ monomer, as evidenced by the RMSD of a complete monomer upon superposition of either CTD or NTD. We also compared the domain conformations between filament and crystal structures by superimposing the NTD of one SmFtsZ monomer (curved state 1) of the filament with NTD of SaFtsZ (PDB 7OHK) and calculated the RMSD for the CTD. We observed that the CTD of SmFtsZ filament is moved away from the NTD resulting in a wider inter-domain cleft between NTD and CTD compared to the crystal structure (Fig. S6B).

We performed a comparative structural analysis of curved state 1 and 2 to elucidate the potential residues involved in inter-monomer bending of FtsZ filament. Consider a FtsZ filament segment consisting of two monomers stacked one over the other (Fig. S7). We first identified the regions in the NTD in the bottom subunit (zone 1 to zone 4) which interacts with CTD in the top subunit (zone 5 and zone 6). Subunits B and C corresponding to those of highest resolution in the two states (Fig. 2A, S3A) were chosen for the analysis. CTDs of a corresponding subunit from the two different states (labelled as subunit ‘0’; Fig. 3A) were superimposed to analyze the change in the NTD movement of the bottom subunit (labelled as ‘-1’). C^α^-C^α^ distances of equivalent residues were compared between the different structural states (Fig. 3A; left). Similarly, CTD movement was tracked by analysing the change in the top subunit (called +1) by superimposing the NTDs of middle subunits from two different states (Fig. 3B; left). The C^α^-C^α^ distances were compared across zone 1 to zone 4 of NTD and zone 5 and zone 6 of CTD to identify the maximum and minimum deviations (Fig. 3A, 3B; right). The C^α^-C^α^ distances were found to be minimum in zone 2 and zone 4 from NTD and zone 5 from CTD suggesting minimal movement in these regions (static zone) during bending of filament and the presence of a hinge or pivot points in these regions (Fig. 3A, B right and supplementary movie 4). In contrast, zone 3 and zone 6 move the maximum (dynamic zones) indicating this interacting surface located opposite to the hinge contains the flexible residues.

**Figure 3.**
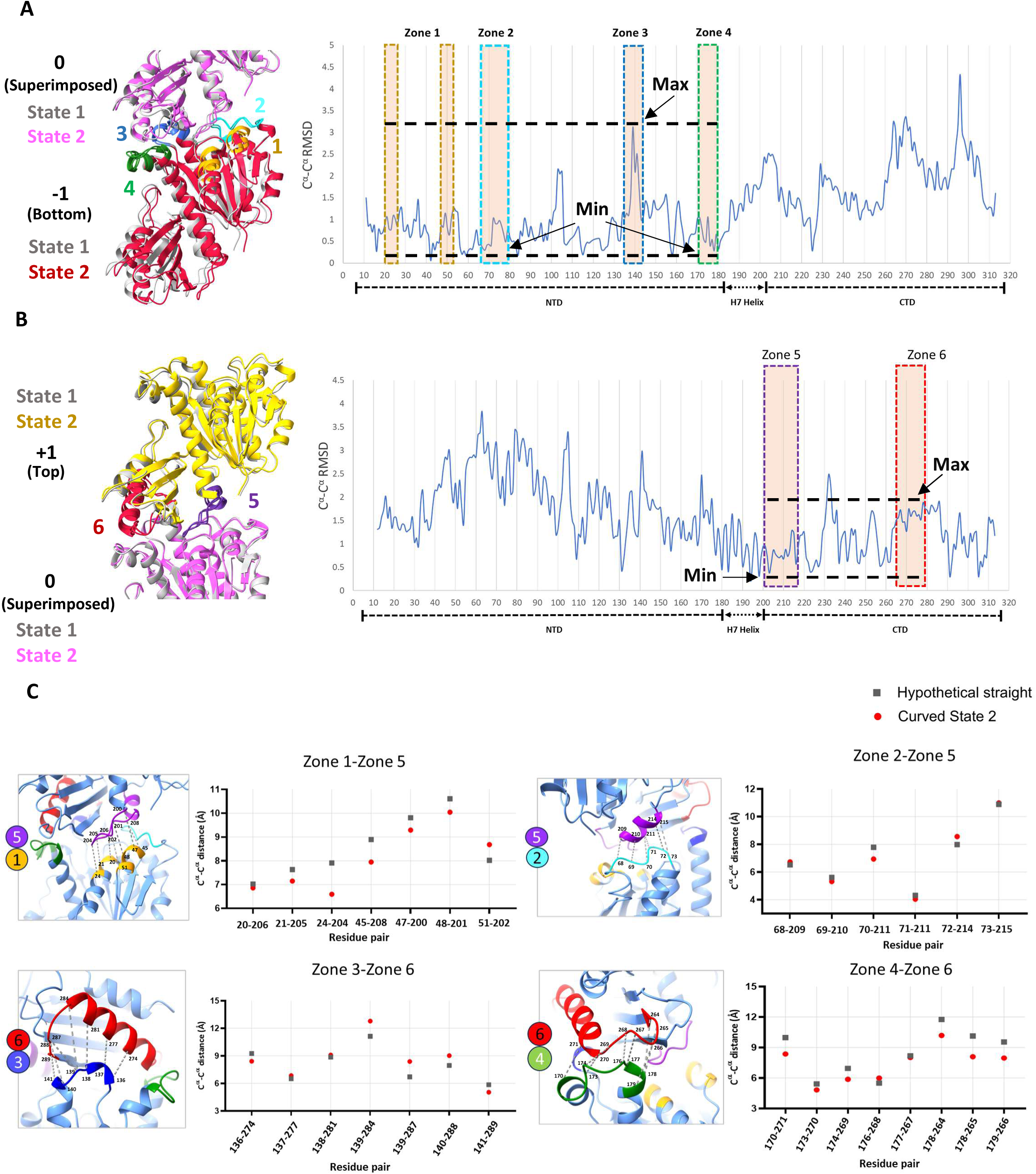
Inter-subunit movements in the SmFtsZ filament at the longitudinal interface determine filament bending. A. Measurements of NTD movements and comparison between state 1 and state 2 structures. Dashed lines are drawn to mark the maximum and minimum deviations across different zones. B. Measurements of CTD movements and comparison between state 1 and state 2 structures. Dashed lines are drawn to mark the maximum and minimum deviations across different zones. C. Measurements of C^α^-C^α^ distances for different interfacing zones at the filament longitudinal interface and comparison between curved state 2 and a hypothetical straight filament. The interfacing residue pairs have been shown in the left figure for respective zones.

To further evaluate the role of specific residues in curvature generation, we measured the C^α^-C^α^ distances of the residues located at the interface zones in two extreme curvature states i.e. the curved state 2 was compared with the corresponding distances in a hypothetical straight filament. The hypothetical straight filament was obtained by fitting SmFtsZ monomer in a straight filament map of KpFtsZ (EMD-60837). We hypothesised that the C^α^-C^α^ distances for the residues located at the hinge region would remain constant in both straight and curved structures whereas residues located at the convex and the concave sides show increased and decreased distances respectively in the curved state compared to the straight one. As expected, the interface residue distances in the zone 1 to zone 5 and zone 4 to zone 6 on the concave side of the curved filaments decreased while zone 2 to zone 5 remained unchanged. Increase in the interface distances was only observed in the zone 3 to zone 6 (residue 139-140 stretch) on the convex side of the filament (Fig. 3C).

### Molecular dynamics simulations support an intrinsic tendency of SmFtsZ protofilaments to bend

To investigate the conformational dynamics of the SmFtsZ protofilament, we performed all-atom molecular dynamics (MD) simulations of a tetramer extracted from curved state 1. The tetramer was solvated in a rhombic dodecahedral water box under periodic boundary conditions (Fig. 4A). Three independent microsecond-long simulations were performed and analysed. To quantify filament bending, we first defined a reference coordinate system using the hypothetical straight SmFtsZ protofilament, with the filament axis aligned along the Z-axis (Fig. 4B). To define the radial direction relative to a microtubule-like geometry, three adjacent protofilaments from a microtubule structure (PDB 3J6E) were aligned to the same reference, and the Y-axis was oriented from the microtubule axis towards the central protofilament. For each MD trajectory, the bottom subunit of the SmFtsZ tetramer was aligned to the corresponding subunit of the straight reference structure. The centre-of-mass position of the top monomer of the tetramer was then projected onto the XY plane, reflecting the tetramer curvature during the simulations (Fig. 4C). Each point in this plot represents one conformation sampled every 1 ns during the last 500 ns of each trajectory. The experimentally determined curved states 1 and 2 were projected onto the same coordinate system for comparison (black and gray dots). In all three trajectories, the protofilaments bent in the direction corresponding to the microtubule lumen (Fig. 4C). The simulated ensemble included conformations comparable in curvature to state 2, although state 2 itself was located at the edge of the sampled distribution. Thus, the MD simulations indicate that state 2 represents one of several accessible curved conformations rather than a uniquely preferred structure.

**Figure 4.**
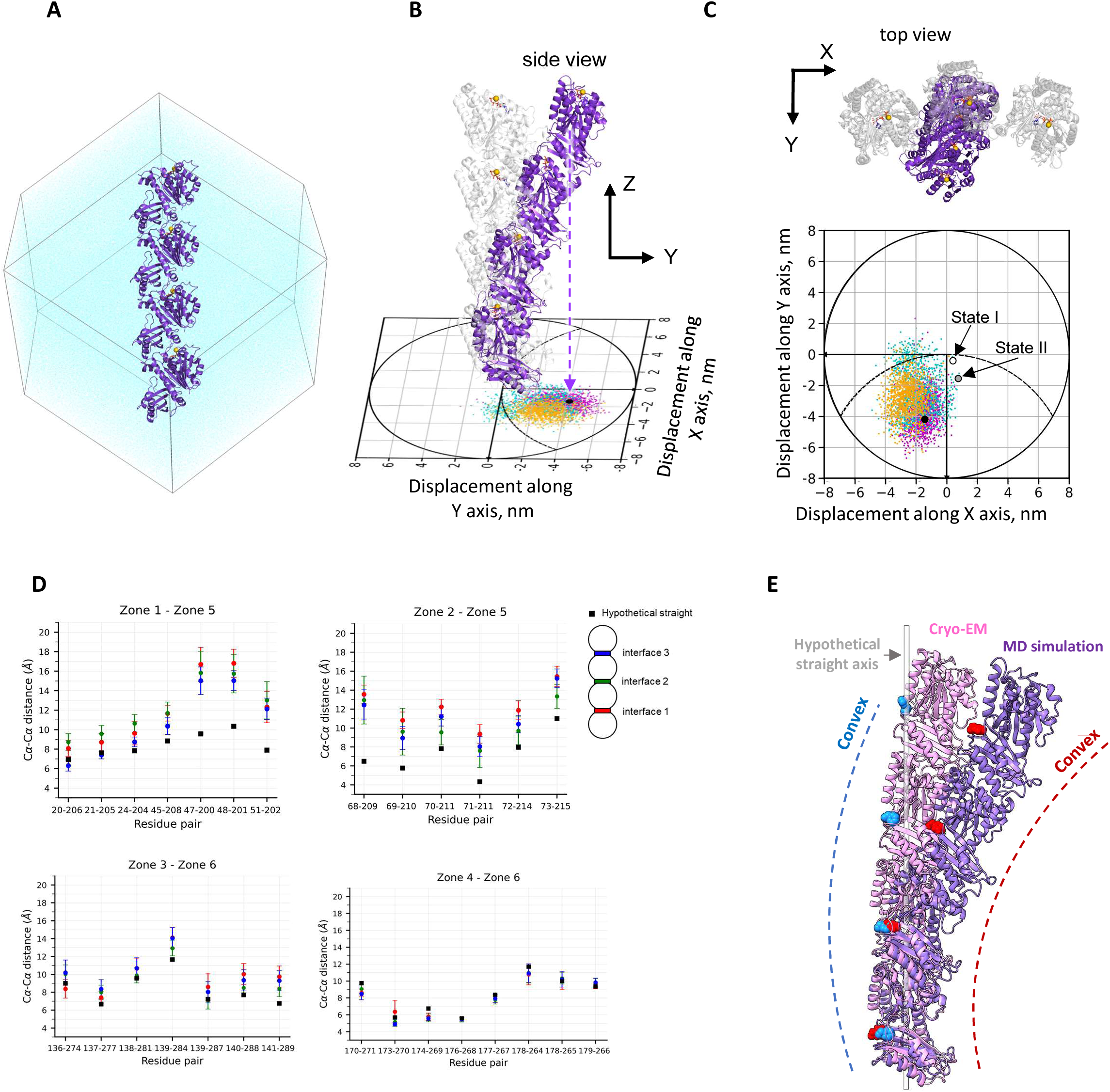
Molecular dynamics of the SmFtsZ filament. A. FtsZ tetramer in a rhombic dodecahedral simulation box. The protein is shown in purple, water molecules in cyan, and Mg²□ ions as gold spheres. B. Gray: reference straight structure of an SmFtsZ tetramer aligned along the Z axis. Purple: a curved conformation sampled during molecular dynamics simulation, with the bottom subunit aligned to the corresponding subunit of the straight reference structure. The center of mass of the top FtsZ subunit is projected onto the XY plane (black dot). C. Gray: three adjacent straight tubulin protofilaments from the microtubule structure (PDB 3J6E), viewed from above; the central protofilament is aligned with the reference straight FtsZ structure. Purple: the same curved FtsZ conformation as in panel B, viewed from above. The lower plot shows projections of the center of mass of the top FtsZ subunit from three independent MD simulations (three colored point clouds), sampled every 1 ns of the simulation. The dashed curve indicates the approximate circumference of a microtubule. The black point marks the central structure of the simulation dataset shown in cyan. For convenience, analogous projections are visualized for experimentally determined FtsZ structures in states 1 and 2. D. Analysis of Cα–Cα distances between residue pairs located at the inter-subunit interfaces in one molecular dynamics simulation. Red, blue, and green points correspond to the three different inter-subunit interfaces; error bars indicate the corresponding fluctuations. Black squares denote the respective distances in the hypothetical straight structure. The schematic on the right indicates the positions of the three inter-subunit interfaces along the simulated FtsZ tetramer. E. Comparison of State 2 structure (pink) with one of the curved states (purple) obtained in simulation run showing the C-terminal end of the filament model (Phe313 atoms marked in sphere style in blue and red color in two structures respectively). White transparent line corresponds to the axis connecting centroids of subunit 1 and 4 of the hypothetical straight filament. Blue and red dotted lines are schematic representations of the monomer C-terminal ends positioning at the convex side of the filaments in cryo -EM and MD simulation respectively.

We next examined the relative motions of the interface regions defined in the previous section. The MD trajectories revealed substantial changes in the relative positions of the residues in zones 1 and 5, as well as in zones 2 and 5, compared with the hypothetical straight protofilament (Fig 4D). In contrast, the relative positions of the residues in zones 3 and 6 and in zones 4 and 6 remained comparatively stable. Structural and dynamic rearrangements in these zones may contribute to the protofilament flexibility. Specifically, compared to the hypothesized straight structure, we see an increase in the distance between contacting residues in one region (zones 1 and 5, 2 and 5), which indicates a decrease in monomer interaction. These structural rearrangements occur due to rotation around a second relatively stable contact area (zones 3 and 6, 4 and 6). Our molecular dynamics simulation trajectories and the curved conformations identified through cryo-EM both indicate the presence of dynamic and stable regions at the monomer-monomer interfaces. Additionally, the consistent orientation of the C-terminal end toward the convex surface of curved filaments in both MD simulations and cryo-EM structures (Fig. 4E) suggests a functional significance of this arrangement which probably marks the membrane side of the filament.

### Lateral interaction straightens FtsZ filament

We next investigated how straight FtsZ filaments are stabilized despite the intrinsic curvature of single protofilaments. Increasing protein concentrations resulted in a significant change in the filament morphology, with the appearance of bundles of straight filaments (Fig. 5A-C). Filament straightening is also observed in SmFtsZ^D211G^ at higher protein concentrations (Fig. 5D-F). Additionally, we observed an increase in the GTPase activity (per µM of FtsZ per minute) with increasing concentration of FtsZ (Fig. 5G). We also performed 2D classification of the double protofilaments which clearly showed absence of any curvature when protofilaments interact laterally (Fig. S8). All 2D class averages of laterally interacting double filaments are straight unlike 2D class averages of single protofilament (Fig. S8, S2). Similar observation was also obtained in earlier reports of laterally interacting straight double filament structure in KpFtsZ bound with ZapA (Fig. 5H) (Fujita et al., 2025).

**Figure 5.**
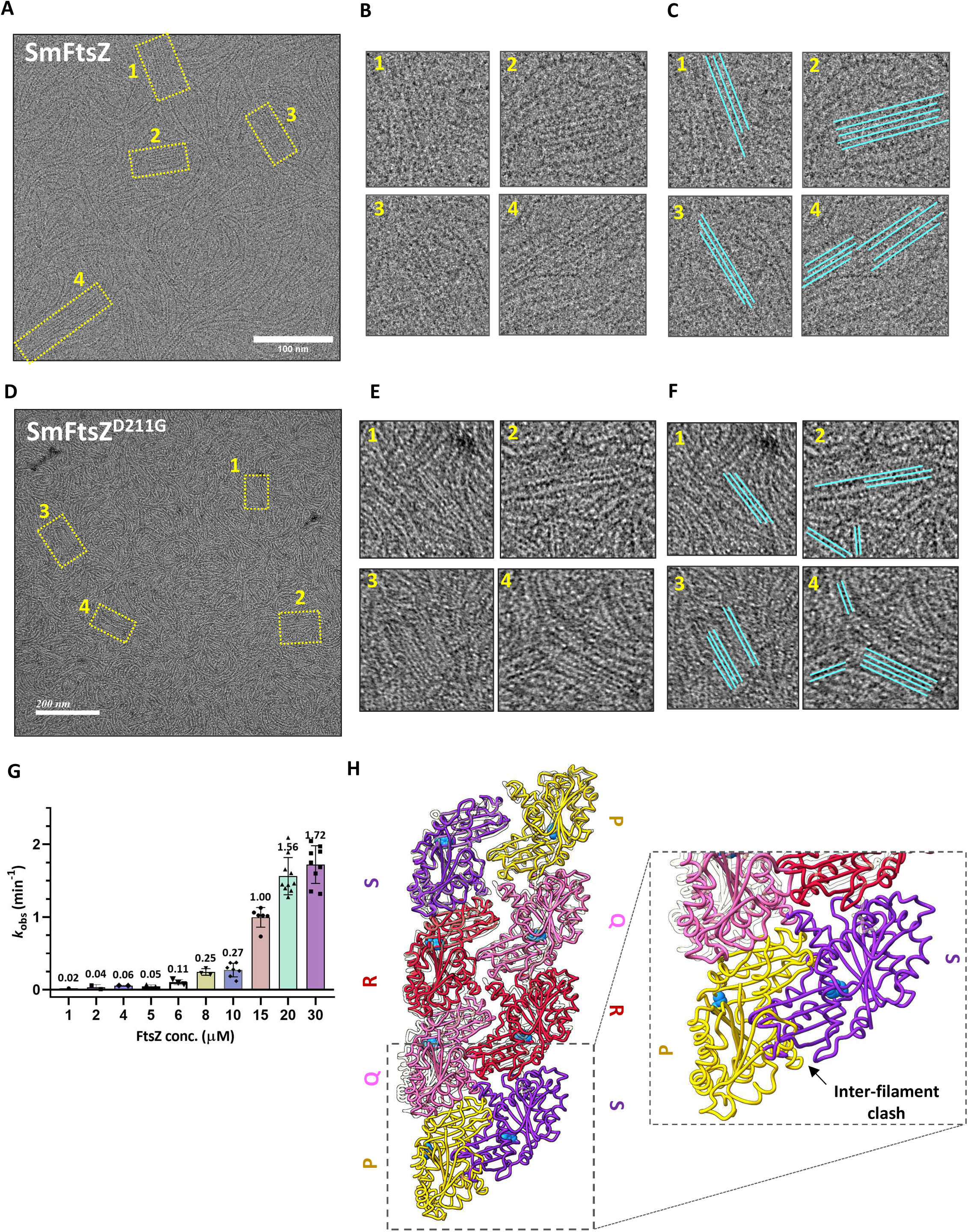
Lateral interactions mediate filament straightening at high concentrations of SmFtsZ. A. Cryo-EM micrographs of SmFtsZ filaments showing bundle formation at higher concentration (i.e. 10 µM). B. Zoomed sections of the four selected regions (yellow boxes) from A. C. Straight filaments are marked with cyan lines in the zoomed images. D. Negative-staining TEM micrographs of SmFtsZ^D211G^ filaments at 1 µM concentration. E. Zoomed sections of the four selected regions (yellow boxes) from D. F. Straight filaments are marked with cyan lines in the zoomed images GTPase assay at increasing concentrations of SmFtsZ. G. GTPase activity of SmFtsZ at various increasing concentrations. H. Superimposition of laterally interacting KpFtsZ-ZapA double protofilament model (PDB-9ISK shown in white transparent color; The chains corresponding to ZapA protein are hidden for better visualization) with curved state 2 protofilaments showing that the curved conformation is unfavorable for lateral interactions due to potential inter-filament clashes. The C-terminal end residue atoms i.e. Phe313 are marked in blue colored sphere style.

Next, we tried to model how the lateral interactions could contribute to straight conformations, by restricting the freedom of movement of the curved protofilaments. We superimposed two protofilaments corresponding to curved state 2 with the previously determined straight antiparallel double protofilament FtsZ structure PDB 9ISK) (Fig. 5H). We observed that laterally interacting curved states are not compatible with the lateral interactions observed in the antiparallel double protofilament interaction in the FtsZ-ZapA complex. Thus, we conclude that the straight FtsZ filaments arise from lateral interactions that suppress the structural plasticity of the interface and hence the intrinsic curvature of individual filaments. High protein concentration or presence of bundling proteins promotes such interactions, effectively overriding the intrinsic bending tendency of individual protofilaments. These findings establish lateral association as a key structural determinant of filament straightening and provide a mechanistic framework for understanding how FtsZ architectures may be regulated *in vivo*.

## Discussion

In eukaryotic cytoskeletal filaments like microtubules, there has been a considerable debate about the relationship between the nucleotide state of the filament and its curvature (reviewed in Gudimchuk and McIntosh, 2021). GTP hydrolysis was initially proposed to directly allosterically alter the conformation of tubulin oligomers, switching them from a straight to a curved state. However, more recent studies indicate that tubulin protofilaments are intrinsically curved irrespective of their bound nucleotide and must be strained to incorporate into the lattice and form lateral bonds. GTP hydrolysis likely weakens these lateral bonds (Manka and Moores, 2018) thereby promoting the release of curved protofilaments from the lattice. The stored strain energy can then be converted into mechanical force (Gudimchuk and Alexandrova, 2023). The prokaryotic tubulin homolog FtsZ is an essential cell division protein that assembles into a dynamic Z-ring at mid-cell and exhibits GTP-dependent treadmilling. Compared to microtubules, FtsZ filaments are shorter, dynamic, and often bundled. All high-resolution structures reported till date for FtsZ filaments have captured straight conformations, obtained upon imposing a helical symmetry with near zero twist in cryo-EM (Fujita et al., 2023; Kumari, Uthaman et al., 2025) or by X-ray crystallography with a filament structure observed in the crystal packing (Ruiz et al., 2023).

While nucleotide hydrolysis–induced curvature is central to microtubule dynamics, whether a similar mechanism is relevant to FtsZ ring constriction remains unknown. Here we show that unconstrained FtsZ protofilaments possess a non-zero intrinsic curvature in the GTP state. Straight filaments emerge from lateral interactions that constrain intrinsically curved protofilaments. Earlier, cryo-EM analysis of KpFtsZ protofilaments have been observed to form straight double filament in the presence of ZapA (Fujita et al., 2025). The intrinsic curvature could be useful for sensing of the membrane curvature such that the filaments are oriented along the cell diameter to aid ring formation at the division site. It is possible that the role of bundling proteins like ZapA, ZapD (and in Spiroplasma, possibly FtsA, SepF) during FtsZ assembly is to align protofilaments and suppress excessive curvature fluctuations, thereby enabling intrinsically curved FtsZ filaments to dynamically shift from curved to straight states based on the occupancy of the interacting partners.

The existence of multiple curved states in a GTP bound form, revealed here with the negative stain EM, high resolution cryo-EM and MD simulations, supports the hypothesis that filament curvature is an inherent property of the FtsZ protofilament, resulting from structural rearrangement of inter-subunit interfaces. Comparison of curved states 1 and 2 primarily reflects variations in the relative orientation of interacting domains, in addition to changes in nucleotide conformation. In both states, we clearly identify the gamma phosphate density in all subunits. Possibly, the slow hydrolysis rate of SmFtsZ enabled us to capture the monomer in a pre-hydrolysis state within the filament. Interestingly, there is a difference in the position of the gamma phosphate in the two states indicating that the pocket allows for sampling various GTP conformations within the curved filaments. We hypothesise that the plasticity of zones 1, 2 and 5 might be restricted in the catalytically optimal geometry of a straight filament. However, the γ-phosphate position in a comparatively higher curvature conformation of the filament (for example, curved state 2) may not be in a compatible geometry for GTP hydrolysis. Hence, the straightening of the filament might lead to stimulation of GTPase activity and efficient treadmilling during ring assembly and constriction. The cause-effect relationship between curvature generation, nucleotide conformation and GTP hydrolysis is unclear due to the limited resolution of the catalytic pocket.

MD simulations confirmed a clear tendency of FtsZ filaments to adopt a curved concave conformation. Notably, none of our simulation runs produced a straight filament conformation. Based on this, we hypothesize that the intrinsically curved conformations represent a relaxed state, while the straight conformation corresponds to a tensed state. The observed interface dynamics from the MD simulations and the curved structures captured by cryo-EM underscore the structural flexibility at the subunit interfaces within single FtsZ protofilaments. The simulated filaments sampled a broad range of curvatures. This flexibility is also evident from the wide scatter in filament radii measured by negative-stain electron microscopy (Fig. 1A, B) and from the conformational heterogeneity revealed by 3D variability analysis of the cryo-electron microscopy structures (Supplementary Movies 1 and 2). The direction of the curvature with a bending of NTD in the concave side in a native GTP bound state of a FtsZ filament supports one of the earlier experimental observations by Osawa et al., 2009. In their liposome reconstitution experiment, they observed a change in curvature of liposomes depending on the presence of membrane targeting site (MTS) at the N- or C-terminal side. A concave depression was observed when the MTS is at the C-terminal end while a convex bulge was observed for an N-terminal MTS. Based on these observations they proposed that the FtsZ protofilament has a preferential bend with C-terminal on the convex side and N-terminal on the concave side. Based on our experimental findings, we propose a model of FtsZ filament assembly, as illustrated in Figure 6. At the onset of cell division, FtsZ filaments are inherently curved which is possibly required to align itself with the short axis of the cell in a similar mechanism as proposed for FtsA by Nierhaus et al., 2022. The curved filaments adopt a GTP conformation that has lower catalytic efficiency. Later, during the Z-ring assembly, presence of accessory proteins (such as FtsA, SepF, ZapA) activate FtsZ by straightening of the filament structure and promoting bundling that results in a condensed ring structure. This conformational change in the filament geometry triggers a conformational change in the catalytic site, resulting in a more productive and efficient hydrolysis necessary for treadmilling.

**Figure 6.**
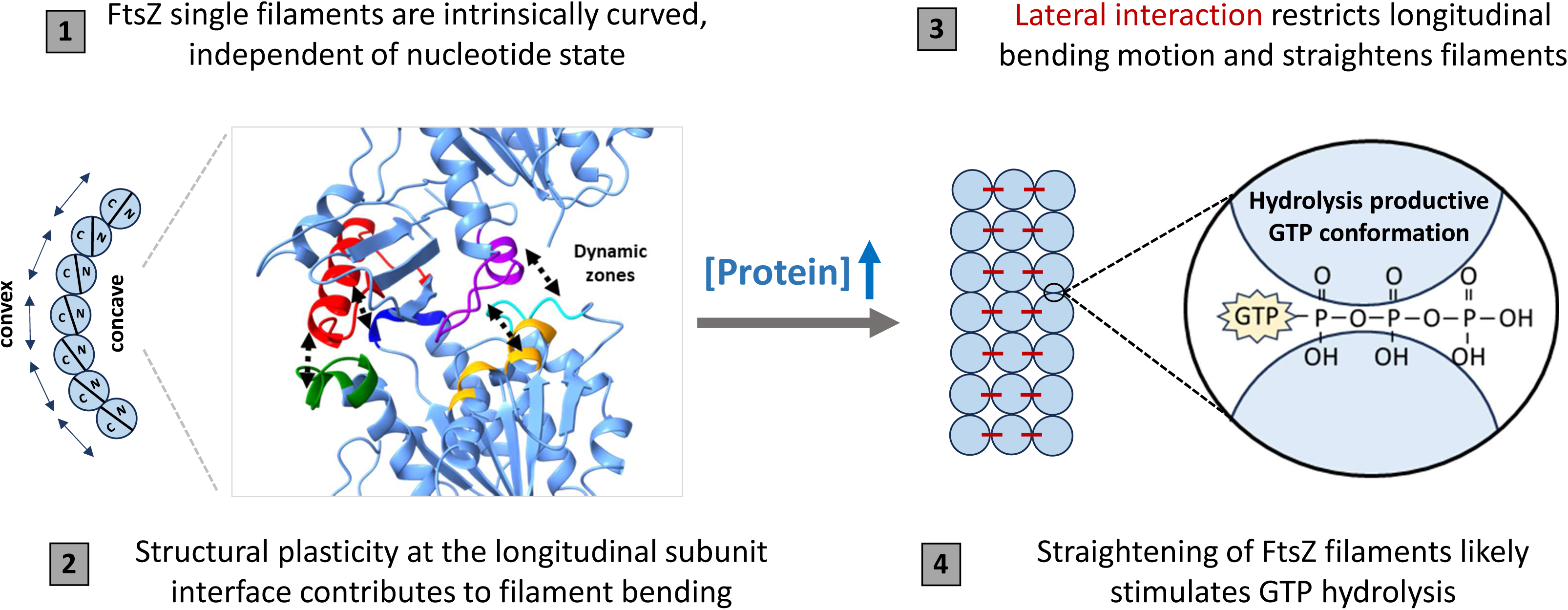
Model of filament bending and straightening in SmFtsZ. (Left) Bent filament is catalytically less efficient and bending is facilitated by dynamic and static interfacing zones at the longitudinal subunit interface. (Right) On the other hand, straight filament is more efficient for GTP hydrolysis. Lateral interactions between protofilaments can be induced by higher FtsZ protein concentration or presence of FtsZ associated proteins.

The present analysis provides mechanistic insight into how bending is achieved in the FtsZ filament which revealed an interplay between a structurally rigid hinge region and a dynamically mobile interface. Notably, these dynamic zones support a lever-like mechanism in which bending is driven by coordinated expansion and contraction of two opposite interfaces. Residues within hinge regions are likely critical for maintaining structural rigidity while those in the dynamic zones may contribute to filament plasticity. Future studies focusing on these residues will be essential to directly link molecular interactions to force generation in FtsZ-mediated cell division.

Together, these findings provide a structural framework for understanding curvature dependence of FtsZ filament function and provide a basis for future studies to evaluate the physiological relevance of the curvature in driving force generation during cell constriction.

## Material and Methods

### Cloning and protein purification

Cloning and purification of SmFtsZ was performed followed by our earlier published protocol in Dutta et al. 2025. Briefly, the construct was expressed in *E. coli* BL21-AI cells by 0.2% arabinose induction at OD_600_= 0.6-0.8 and was grown at 30 °C for 5 hours. Cells were harvested by centrifugation at 4,000 xg at 4 °C for 20□mins and the cell pellet was flash-frozen in liquid nitrogen and stored at -80 °C until processed. The cell pellet was thawed in ice for 10 mins and then resuspended in a lysis buffer (50□mM Tris-HCl pH 8.0, 50□mM NaCl, 10% glycerol) followed by sonication to lyse the cells. The total lysate was spun at 30,000 xg for 1 hour to remove the cell debris and the clear supernatant was loaded into HiTrap Q HP 5□ml column (Cytiva) equilibrated in Buffer A50 (50□mM Tris-HCl pH 8.0, 50□mM NaCl). Protein was eluted in 5 mL fractions using a linear NaCl gradient from 50 mM to 400□mM over 40 column volumes. Peak fractions containing proteins were confirmed from SDS PAGE, collected and pooled together for dialysis against A50 buffer and loaded into 10/100 GL MonoQ column (Cytiva) to improve purity. Finally, the pure fractions were confirmed from SDS PAGE gel and pooled for dialysis in the HK50 buffer (50□mM HEPES pH 7.4, 50□mM KCl). After dialyzing for 1.5 hours, the protein was concentrated using centricon (with 10 kDa membrane cut off), flash-frozen in aliquots and stored at -80 °C. The cloning and purification of SmFtsZ^D211G^ was performed as per the earlier published protocol in Chakraborty et al., 2024.

All purified proteins were run on 12% SDS-PAGE for checking the purity level and quality (Figure S9).

### Negative staining sample preparation for transmission electron microscopy (TEM)

Polymerization reaction was set up in HK50 buffer with specific concentration of proteins, 5□mM MgCl_2_ and 2 mM GTP and incubated for 10 min at 30 °C. For visualizing under TEM, 5 μL of reaction was loaded on a glow discharged (15 mA for 25 sec, negative surface) carbon coated copper grid and incubated for 2 minutes before blotting out excess sample from the side. Then 5□μL 1.5% (w/v) uranyl acetate stain solution was added on the grid and blotted using Whatman filter paper from the side after 30 seconds. Grid was air-dried and imaging was performed in TEM (model: JEM-2200FS Jeol Ltd.) operated at 200□kV.

### Filament curvature analysis of TEM micrographs

Curvature of the filaments were measured by manually tracking filaments in TEM micrographs using kappa plugin (Mary and Brouhard, 2019) within Fiji software (Schindelin et al., 2012). All the plots were generated using GraphPad Prism software.

### Cryo-EM sample preparation and data collection

For single filament reconstruction of SmFtsZ, 2 µM protein sample was used in HK50 buffer with 5□mM MgCl_2_ and 2 mM GTP. The reaction mixture was incubated for 10 min at 30 °C before applying on the grids. Quantifoil grids (R1.2/1.3 Au 200 mesh) were glow-discharged at 25 mA for 60 sec. Upon 3 µL of sample application, grid was blotted using Vitrobot Mark IV with a force of 0 and a time of 3.5□s and then immediately flash frozen by plunging into liquid ethane. Cryo-EM image datasets were acquired using Titan Krios G3i microscope (Thermo Fisher Scientific, USA) operated at 300□kV using a Falcon 3 detector in counting mode. The defocus range was set between -1.8□μm and -3.0□μm. The nominal magnification was 75,000×, corresponding to 1.07□Å pixel size. Each movie was fractionated into 25 frames with a total dose of 25.67□e/Å^2^.

For double protofilament reconstruction, a higher concentration i.e. 10 µM of SmFtsZ protein was used in HK50 buffer with 5□mM MgCl_2_ and 2 mM GTP. The reaction mixture was incubated for 10 min at 30 °C before applying on the grids. Imaging was performed with a magnification of 59,000×, corresponding to 1.38□Å pixel size. Each movie was fractionated into 16 frames with a total dose of 17.74□e/Å^2^. Grids and glow discharge and freezing condition and other microscope parameters were the same as mentioned for the previous data set.

### Cryo-EM image processing, model building and structure analysis

Cryo-EM data processing was done in cryoSPARC (Punjani et al., 2017) version 4.7.1. A total of 1,080 movies were imported and motion corrected, and the contrast transfer functions (CTFs) were estimated. 965 micrographs with CTF resolutions above 5□Å were selected to process further for particle picking. Initially particle picking was done using template free filament tracer with a filament diameter of 40 Å. After running a 2D classification step, best classes were chosen for template-based filament picking. In the filament tracer job, separation distance was kept as 0.5 and hysteresis threshold (min, max) was optimized as (85, 90) to increase the number of particles. All the other parameters were set as default. Next, the particles were extracted with a box size of 320 pixels. After two rounds of 2D classification, best 2D class averages containing 387,392 particles with features of secondary structure (resolution better than 8 Å) were selected for an ab-initio 3D reconstruction (3 classes). After heterogenous refinement, straight and curved classes were selected for non-uniform refinement (Punjani et al., 2020) followed by global and local CTF refinements. A final step of non-uniform refinement followed by a local refinement step which yielded an overall map resolution of 3.4 Å and 3.9 Å for curved state 1 and state 2 respectively. 3D variability analysis (Punjani et al., 2021) with the final set of particles was performed in cryoSPARC.

The initial model of SmFtsZ T state was generated in SWISS-MODEL (Waterhouse et al., 2020) using the crystal structure of *Staphylococcus aureus* FtsZ as template (PDB 3VOA). The T state SmFtsZ model was used for further model building using COOT ver. 0.9.8.7 (Emsley et al., 2010). During model building, four monomers and GTP were fitted into the electron density map to generate the filament model of SmFtsZ which was then real space refined in PHENIX ver. 1.21.2 (Adams, P. D. et al., 2010). ChimeraX ver. 1.10.1 was used for all structure related measurements and making figures. Plots were generated using Microsoft Excel and GraphPad Prism software.

### GTPase activity measurement

The GTPase activity was measured using Malachite green assay (Feng et al. 2011). The protein was spun at 21,000xg at 4°C for 10 mins and 20 μL reaction was set with the required concentration of protein in the reaction buffer (50 mM HEPES pH 7.4, 50 mM KCl), 2mM of GTP, and 5 mM MgCl_2_. Then this reaction mixture was incubated at 30°C for 10 minutes. To stop GTPase reaction, 5 μL of 0.5M EDTA was added in 20 μL reaction. 100 μL of the malachite green mixture was added into the final 25 μL sample and waited for 10 mins for the color development and absorbance was measured at 630 nm wavelength. Phosphate solutions of known concentrations were used as standards to estimate the released phosphate amount in the reaction. All the plots were generated using GraphPad Prism software.

### Molecular dynamics simulations

Molecular model of the SmFtsZ tetramer was based on the state 1 structure obtained by single particle cryo-EM. Propka (Olsson et al., 2011) was used to calculate the unknown degree of protonation of ionizable amino acid residues and Dowser (Morozenko and Stuchebrukhov, 2016) was used to identify and solvate cavities inside the protein. The SmFtsZ tetramer was placed in a rhombic dodecahedral simulation box with a side length of 21 nm with periodic boundary conditions filled with TIP3P water molecules. The ionic strength of the solution was set at 100 mM by adding K^+^ and Cl^−^ ions so that the total charge of the system was zero. Simulations were performed using the GROMACS 2025 software package, which allows parallel computing on hybrid architecture (Páll et al., 2020) with the CHARMM36m force field (Huang et al., 2017).

We minimized the system energy using the steepest descent algorithm followed by a two-step equilibration: first, we conducted 1 ns long simulation with constrained positions of all heavy protein atoms at constant pressure and temperature; second, we carried out 5 ns long simulation with constrained positions of protein backbone atoms, using the Berendsen barostat (time constant 4.0 fs, compressibility 4.5×10^−5^ bar^−1^) and the Berendsen thermostat. The production simulation runs were carried out in the NPT ensemble at 300 K, using the Parrinello-Rahman algorithm (Parrinello and Rahman, 1981) and the V-rescale thermostat for a duration of 1 μs each. The particle mesh Ewald method was used to treat the long-range electrostatics. Mass rescaling (partial transfer of mass from heavy atoms to bound hydrogens (Feenstra et al., 1999) allowed molecular dynamics simulations with 2 fs time step. MDAnalysis (Michaud-Agrawal et al., 2011) was used for the analysis of molecular simulation data such as C_α_-C_α_ distances of the residue pairs through molecular dynamics trajectories. PyMOL (The PyMOL Molecular Graphics System, Version 3.0 Schrödinger, LLC). Bending of the FtsZ tetramer was analyzed by plotting projections of the center of mass of the top subunit, using a custom Python script, as described previously (Fedorov et al., 2019).

## Author Contributions

SD and PG conceived the project and designed the experiments. SD conducted all experiments and analyzed the data except mentioned otherwise. JC contributed to cloning and purification of SmFtsZ^D211G^ construct. IK, NG and EK designed molecular dynamic simulations and analyzed data. EK performed the simulations. SD and PG wrote the manuscript with inputs from NG. PG supervised the project and acquired funding. NG and IK supervised the computational part of the project and acquired funding. All authors read and approved the final submitted version of the manuscript.

## Acknowledgments

We acknowledge IISER Pune for all the facilities to carry out this research, the National Electron cryoMicroscopy Facility, BLiSC Campus, Bangalore where the cryo-EM data was collected. We thank Dr. Sucharita Bose and Dr. Kutti R Vinothkumar for helping with cryo-EM sample preparation and data collection, Ilya Volkhin and Vladimir Fedorov for assistance with MD simulations and useful discussions. We acknowledge Pune Biotech Cluster, Department of Biotechnology (DBT) India for providing computational facility and Transmission Electron Microscopy facility at the Department of Physics, IISER Pune. We acknowledge fellowships from CSIR, India to SD; IISER Pune to JC. Research work in the lab of PG was supported by the Department of Biotechnology (DBT) Membrane Structural Biology Program grant (BT/PR28833/BRB/10/1705/2018), S. Ramachandran National Bioscience Award Research grant (HRD-20/7/2024-HRD-DBT), Ministry of Education STARS (2023-1015), and IISER Pune. Molecular dynamics simulations were funded by the Russian Science Foundation, research project 25-44-01015.

## References

1. Adams, P.D., Afonine, P.V., Bunkóczi, G., Chen, V.B., Davis, I.W., Echols, N., Headd, J.J., Hung, L.W., Kapral, G.J., Grosse-Kunstleve, R.W. and McCoy, A.J. (2010). PHENIX: a comprehensive Python-based system for macromolecular structure solution. Biological crystallography, 66(2), 213–221.

2. Alushin, G. M., Lander, G. C., Kellogg, E. H., Zhang, R., Baker, D., & Nogales, E. (2014). High-resolution microtubule structures reveal the structural transitions in αβ-tubulin upon GTP hydrolysis. Cell, 157(5), 1117–1129.

3. Bisson-Filho, A.W., Hsu, Y.P., Squyres, G.R., Kuru, E., Wu, F., Jukes, C., Sun, Y., Dekker, C., Holden, S., VanNieuwenhze, M.S. and Brun, Y.V., 2017. Treadmilling by FtsZ filaments drives peptidoglycan synthesis and bacterial cell division. Science, 355(6326), pp.739–743.

4. Brouhard, G. J., & Rice, L. M. (2018). Microtubule dynamics: an interplay of biochemistry and mechanics. Nature reviews Molecular cell biology, 19(7), 451–463.

5. Chakraborty, J., Poddar, S., Dutta, S., Bahulekar, V., Harne, S., Srinivasan, R., & Gayathri, P. (2024). Dynamics of interdomain rotation facilitates FtsZ filament assembly. Journal of Biological Chemistry, 300(6).

6. Corbin, L.C., and Erickson, H.P. (2020). A unified model for treadmilling and nucleation of single-stranded FtsZ protofilaments. Biophys J 119: 792–805.

7. Dajkovic, A., Mukherjee, A., and Lutkenhaus, J. (2008) Investigation of regulation of FtsZ assembly by SulA and development of a model for FtsZ polymerization. J Bacteriol 190: 2513–2526.

8. Deng, Y., Sun, M., & Shaevitz, J. W. (2011). Direct measurement of cell wall stress stiffening and turgor pressure in live bacterial cells. Physical review letters, 107(15), 158101.

9. Desai, A., & Mitchison, T. J. (1997). Microtubule polymerization dynamics. Annual review of cell and developmental biology, 13(1), 83–117.

10. Du, S., Pichoff, S., Kruse, K., and Lutkenhaus, J. (2018) FtsZ filaments have the opposite kinetic polarity of microtubules. Proc Natl Acad Sci 115: 10768–10773.

11. Dutta, S., Poddar, S., Chakraborty, J., Srinivasan, R. and Gayathri, P., 2025. Membrane binding and cholesterol sensing motif in Mycoplasma genitalium FtsZ: a novel mode of membrane recruitment for bacterial FtsZ. Biochemistry, 64(8), pp.1864–1877.

12. Emsley, P., Lohkamp, B., Scott, W. G. & Cowtan, K. (2010). Features and development of Coot. Acta Crystallogr. D. Biol. Crystallogr. 66, 486–501.

13. Erickson, H. P., & Osawa, M. (2017). FtsZ constriction force–curved protofilaments bending membranes. Prokaryotic Cytoskeletons: Filamentous Protein Polymers Active in the Cytoplasm of Bacterial and Archaeal Cells, 139–160.

14. Fedorov, V.A., Orekhov, P.S., Kholina, E.G., Zhmurov, A.A., Ataullakhanov, F.I., Kovalenko, I.B., and Gudimchuk, N.B. (2019). Mechanical properties of tubulin intra- and inter-dimer interfaces and their implications for microtubule dynamic instability. PLoS Comput Biol 15(8), e1007327.

15. Feenstra KA, Hess B, Berendsen HJC. Improving efficiency of large time-scale molecular dynamics simulations of hydrogen-rich systems. J Comput Chem. 1999; 20: 786–798.

16. Fujita, J., Harada, R., Maeda, Y., Saito, Y., Mizohata, E., Inoue, T., Shigeta, Y. and Matsumura, H. (2017). Identification of the key interactions in structural transition pathway of FtsZ from Staphylococcus aureus. Journal of Structural Biology, 198(2), 65–73.

17. Fujita, J., Amesaka, H., Yoshizawa, T., Hibino, K., Kamimura, N., Kuroda, N., Konishi, T., Kato, Y., Hara, M., Inoue, T. and Namba, K., 2023. Structures of a FtsZ single protofilament and a double-helical tube in complex with a monobody. Nature Communications, 14(1), p.4073.

18. Fujita, J., Kasai, K., Hibino, K., Kagoshima, G., Kamimura, N., Tobita, S., Kato, Y., Uehara, R., Namba, K., Uchihashi, T. and Matsumura, H., 2025. Structural basis for the interaction between the bacterial cell division proteins FtsZ and ZapA. Nature Communications, 16(1), p.5985.

19. Gudimchuk, N.B. and McIntosh, J.R., 2021. Regulation of microtubule dynamics, mechanics and function through the growing tip. Nature reviews Molecular cell biology, 22(12), pp.777–795.

20. Gudimchuk, N.B. and Alexandrova, V.V., 2023. Measuring and modeling forces generated by microtubules. Biophysical Reviews, 15(5), pp.1095–1110.

21. Hsin, J., Gopinathan, A. and Huang, K.C., 2012. Nucleotide-dependent conformations of FtsZ dimers and force generation observed through molecular dynamics simulations. Proceedings of the National Academy of Sciences, 109(24), pp.9432–9437.

22. Huang J, Rauscher S, Nawrocki G, Ran T, Feig M, de Groot BL, Grubmüller H, MacKerell AD Jr. CHARMM36m: an improved force field for folded and intrinsically disordered proteins. Nat Methods. 2017 Jan;14(1):71–73.

23. Huecas, S., Llorca, O., Boskovic, J., Martín-Benito, J., Valpuesta, J.M., and Andreu, J.M. (2008). Energetics and geometry of FtsZ Polymers: Nucleated self-Assembly of single protofilaments. Biophys J 94: 1796–1806.

24. Knossow, M., Campanacci, V., Khodja, L.A., and Gigant, B. (2020) The mechanism of tubulin assembly into microtubules: Insights from structural studies. iScience 23: 101511.

25. Kumari, J., Uthaman, A., Bose, S., Kundu, A., Sharma, V., Dutta, S., Dhar, A., Roy, S., Srinivasan, R., Pande, S. and Vinothkumar, K.R., 2025. Distinct filament morphology and membrane tethering features of the dual FtsZ paralogs in Odinarchaeota. The EMBO Journal, 44(21), p.5940.

26. Lan, G., Daniels, B.R., Dobrowsky, T.M., Wirtz, D. and Sun, S.X., 2009. Condensation of FtsZ filaments can drive bacterial cell division. Proceedings of the National Academy of Sciences, 106(1), pp.121–126.

27. Li, Y., Hsin, J., Zhao, L., Cheng, Y., Shang, W., Huang, K.C., Wang, H.W. and Ye, S. (2013). FtsZ protofilaments use a hinge-opening mechanism for constrictive force generation. Science, 341(6144), pp.392–395.

28. Lu, C., Stricker, J., & Erickson, H. P. (2001). Site-specific mutations of FtsZ-effects on GTPase and in vitro assembly. BMC microbiology, 1(1), 7.

29. Mandelkow, E. M., Mandelkow, E., & Milligan, R. A. (1991). Microtubule dynamics and microtubule caps: a time-resolved cryo-electron microscopy study. The Journal of cell biology, 114(5), 977–991.

30. Manka, S.W. and Moores, C.A. (2018). The role of tubulin–tubulin lattice contacts in the mechanism of microtubule dynamic instability. Nature structural & molecular biology, 25(7), pp.607–615.

31. Mary, H. and Brouhard, G.J., 2019. Kappa (κ): analysis of curvature in biological image data using B-splines. BioRxiv, p.852772.

32. Meng EC, Goddard TD, Pettersen EF, Couch GS, Pearson ZJ, Morris JH, Ferrin TE. (2023) UCSF ChimeraX: Tools for structure building and analysis. Protein Sci. 2023 Nov;32(11):e4792.

33. Merino-Salomón, A., Schneider, J., Babl, L., Krohn, J.H., Sobrinos-Sanguino, M., Schäfer, T., Luque-Ortega, J.R., Alfonso, C., Jiménez, M., Jasnin, M. and Schwille, P. (2023). Crosslinking by ZapD drives the assembly of short FtsZ filaments into toroidal structures in solution. eLife13:RP95557.

34. Michaud-Agrawal N., Denning E. J., Woolf T. B., and Beckstein O. MDAnalysis: A Toolkit for the Analysis of Molecular Dynamics Simulations. J. Comput. Chem. 32 (2011), 2319–2327.

35. Matsui, T., Han, X., Yu, J., Yao, M., & Tanaka, I. (2014). Structural change in FtsZ induced by intermolecular interactions between bound GTP and the T7 loop. Journal of Biological Chemistry, 289(6), 3501–3509.

36. Morozenko, A. and Stuchebrukhov, A.A., (2016). Dowser++, a new method of hydrating protein structures. Proteins: Structure, Function, and Bioinformatics, 84(10), pp.1347–1357.

37. Nierhaus, T., McLaughlin, S.H., Bürmann, F., Kureisaite-Ciziene, D., Maslen, S.L., Skehel, J.M., Yu, C.W., Freund, S.M., Funke, L.F., Chin, J.W. and Löwe, J., 2022. Bacterial divisome protein FtsA forms curved antiparallel double filaments when binding to FtsN. Nature microbiology, 7(10), pp.1686–1701.

38. Nguyen, L.T., Oikonomou, C.M. and Jensen, G.J., (2021). Simulations of proposed mechanisms of FtsZ-driven cell constriction. Journal of Bacteriology, 203(3), pp.10–1128.

39. Olsson MHM, Søndergaard CR, Rostkowski M, Jensen JH. (2011). PROPKA3: Consistent Treatment of Internal and Surface Residues in Empirical pKa Predictions. J Chem Theory Comput.; 7: 525–537.

40. Osawa, M. and Erickson, H.P. (2013). Liposome division by a simple bacterial division machinery. Proceedings of the National Academy of Sciences, 110(27), pp.11000–11004.

41. Parrinello M, Rahman A. (1981). Polymorphic transitions in single crystals: A new molecular dynamics method. J Appl Phys.; 52: 7182–7190.

42. Punjani A, Rubinstein JL, Fleet DJ, Brubaker MA (2017) cryoSPARC: algorithms for rapid unsupervised cryo-EM structure determination. Nat Methods 14:290–296.

43. Punjani, A., Zhang, H. & Fleet, D.J. (2020). Non-uniform refinement: adaptive regularization improves single-particle cryo-EM reconstruction. Nat Methods 17, 1214–1221.

44. Punjani, A. & Fleet, D.J. 3D variability analysis: Resolving continuous flexibility and discrete heterogeneity from single particle cryo-EM. Journal of Structural Biology, Volume 213, Issue 2, 2021.

45. Ramirez-Diaz, D. A., Merino-Salomón, A., Meyer, F., Heymann, M., Rivas, G., Bramkamp, M., & Schwille, P. (2021). FtsZ induces membrane deformations via torsional stress upon GTP hydrolysis. Nature communications, 12(1), 3310.

46. Reynolds, M.J., Hachicho, C., Carl, A.G., Gong, R. and Alushin, G.M. (2022). Bending forces and nucleotide state jointly regulate F-actin structure. Nature, 611(7935), pp.380–386.

47. Ruiz, F. M., Huecas, S., Santos-Aledo, A., Prim, E. A., Andreu, J. M., & Fernández-Tornero, C. (2022). FtsZ filament structures in different nucleotide states reveal the mechanism of assembly dynamics. PLoS biology, 20(3), e3001497.

48. Schindelin, J., Arganda-Carreras, I., Frise, E., Kaynig, V., Longair, M., Pietzsch, T., Preibisch, S., Rueden, C., Saalfeld, S., Schmid, B. and Tinevez, J.Y., 2012. Fiji: an open-source platform for biological-image analysis. Nature methods, 9(7), pp.676–682.

49. Squyres, G. R., Holmes, M. J., Barger, S. R., Pennycook, B. R., Ryan, J., Yan, V. T., & Garner, E. C. (2021). Single-molecule imaging reveals that Z-ring condensation is essential for cell division in Bacillus subtilis. Nature microbiology, 6(5), 553–562.

50. Szilárd Páll, Artem Zhmurov, Paul Bauer, Mark Abraham, Magnus Lundborg, Alan Gray, Berk Hess, Erik Lindahl; Heterogeneous parallelization and acceleration of molecular dynamics simulations in GROMACS. J. Chem. Phys. 7 October 2020; 153 (13): 134110.

51. Szwedziak, P., & Ghosal, D. (2017). FtsZ-ring architecture and its control by MinCD. Prokaryotic Cytoskeletons: Filamentous Protein Polymers Active in the Cytoplasm of Bacterial and Archaeal Cells, 213–244.

52. Waterhouse A, Bertoni M, Bienert S, Studer G, Tauriello G, Gumienny R, Heer FT, de Beer TAP, Rempfer C, Bordoli L, Lepore R, Schwede T. (2018). SWISS-MODEL: homology modelling of protein structures and complexes. Nucleic Acids Res 46, W296–W303.

53. Wagstaff, J.M., Tsim, M., Oliva, M.A., García-Sanchez, A., Kureisaite-Ciziene, D., Andreu, J.M., and Löwe, J. (2017) A polymerization-associated structural switch in FtsZ that enables treadmilling of model filaments. mBio.

54. Wagstaff, J. M., Planelles-Herrero, V. J., Sharov, G., Alnami, A., Kozielski, F., Derivery, E., & Löwe, J. (2023). Diverse cytomotive actins and tubulins share a polymerization switch mechanism conferring robust dynamics. Science advances, 9(13), eadf3021.

55. Whatmore, A. M., & Reed, R. H. (1990). Determination of turgor pressure in Bacillus subtilis: a possible role for K+ in turgor regulation. Microbiology, 136(12), 2521–2526.

56. Whitley, K.D., Jukes, C., Tregidgo, N., Karinou, E., Almada, P., Cesbron, Y., et al. (2021) FtsZ treadmilling is essential for Z-ring condensation and septal constriction initiation in Bacillus subtilis cell division. Nat Commun 12: 2448.

57. Xiao, J., & Goley, E. D. (2016). Redefining the roles of the FtsZ-ring in bacterial cytokinesis. Current opinion in microbiology, 34, 90–96.

58. Yang, X., & Liu, R. (2022). How does FtsZ’s treadmilling help bacterial cells divide?. Biocell, 46(11).

