## Supplementary Table for "Lateral interactions override nucleotide state in determining FtsZ filament curvature"

Figure S1

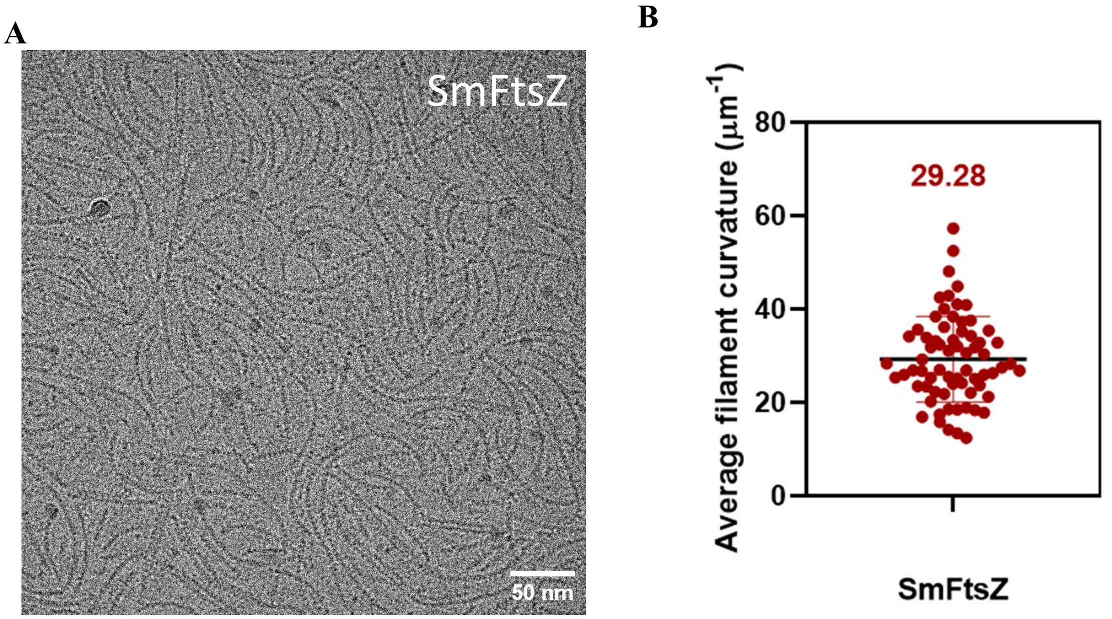


**Figure S1. Filament curvature analysis of SmFtsZ cryo-EM sample**

A. Representative cryo-EM micrographs for SmFtsZ GTP showing similar curvature as negative staining TEM. B. Curvature analysis of SmFtsZ filaments in cryo condition. Mean value is indicated above the data plot.

**Figure S2**

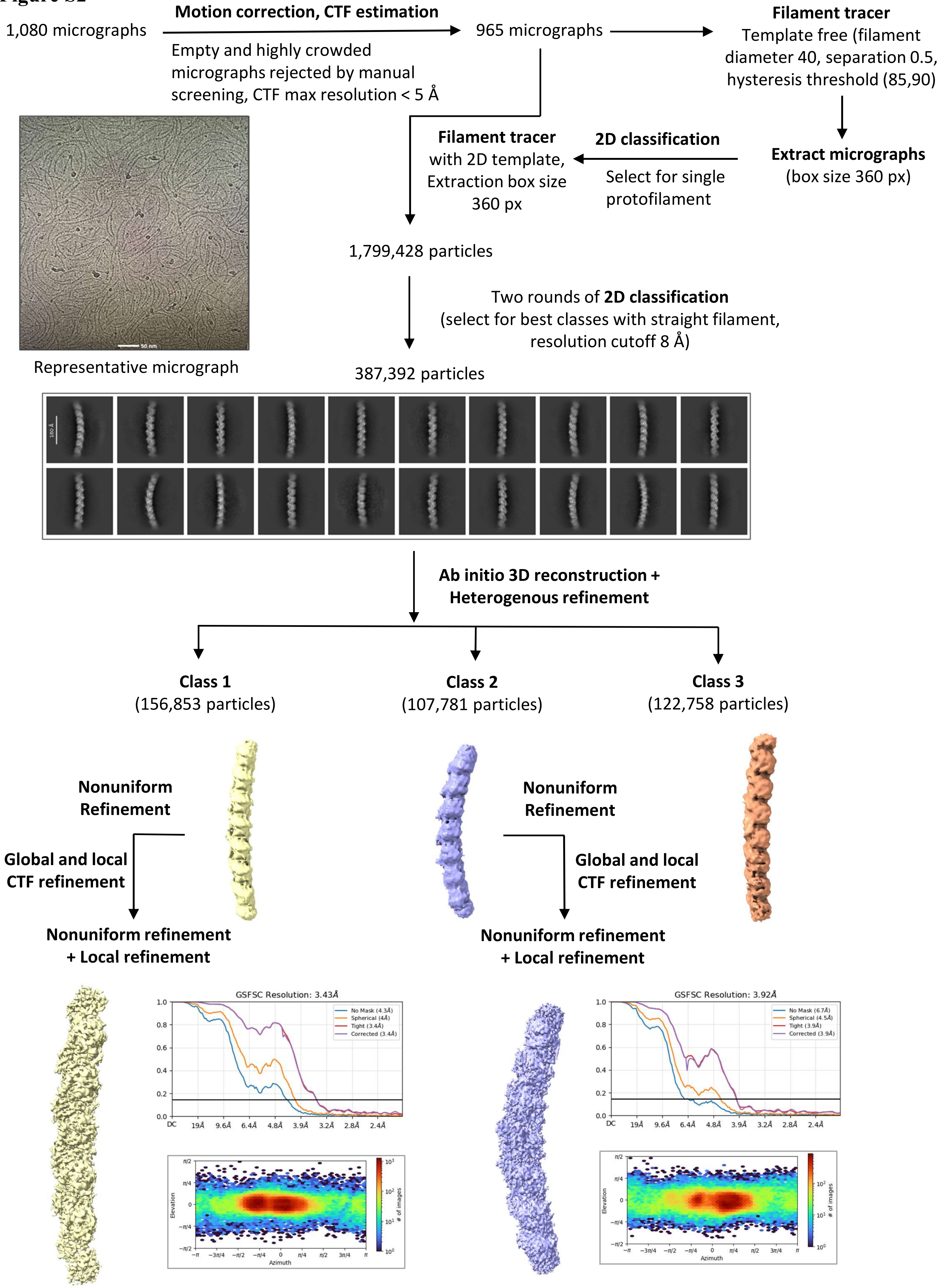


**Figure S2. Cryo-EM data processing workflow for SmFtsZ single filament**

**Figure S3**

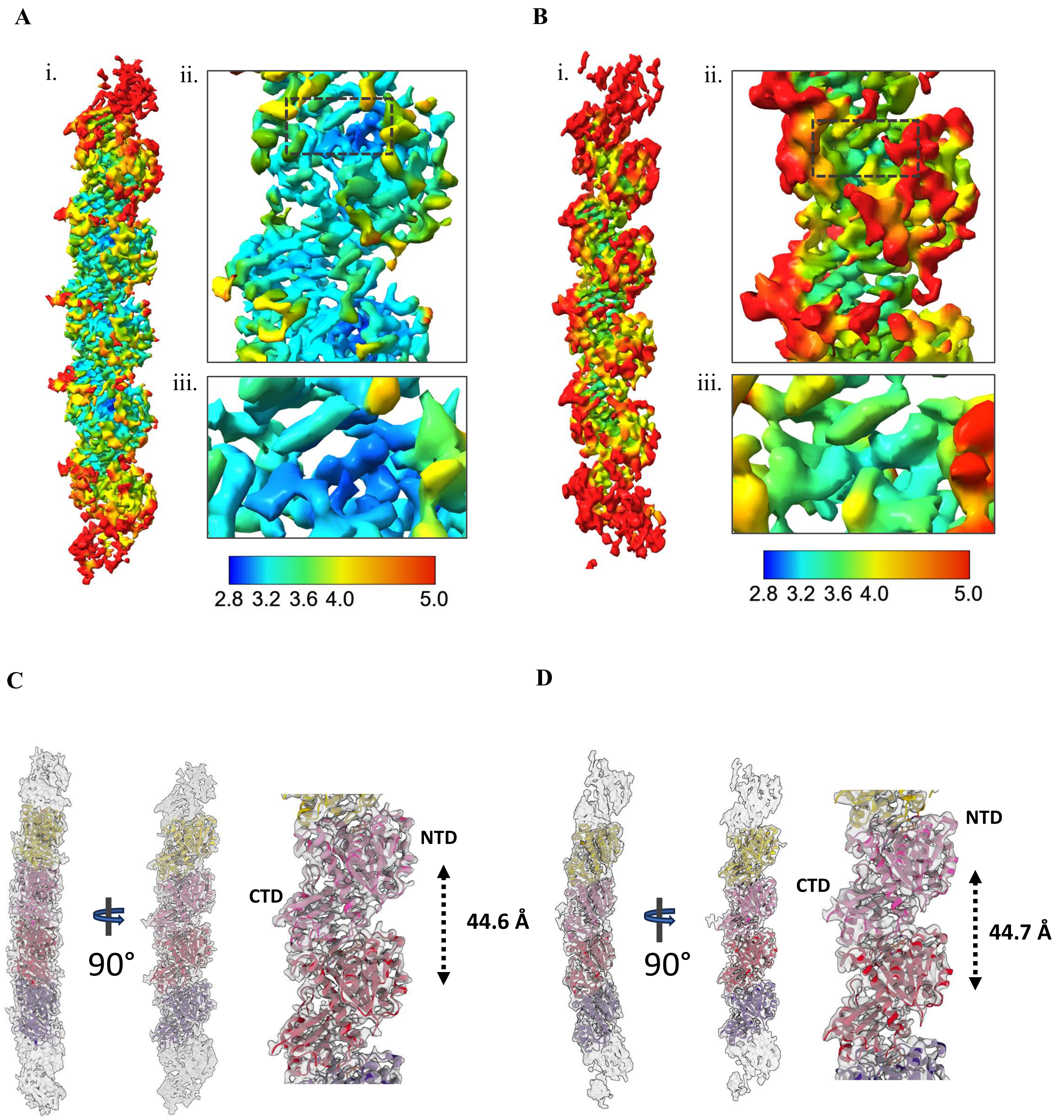


**Figure S3. Cryo-EM analysis SmFtsZ protofilaments in two curvature states**

A, B. The local resolution distributions of state 1 (A) and state 2 (B) maps are colored as in the color bar. All values are reported in Å. Close-up view of single monomer and GTP. C, D. Fitting of atomic filament model in the cryo-EM maps for state 1 (C) and state 2 (D) structures at 0.6 contour level. Average centroid to centroid distances are marked.

**Supplementary Table 1: Refinement statistics of SmFtsZ state 1 and state 2 filament models**

| | **State 1** | **State 2** |
|---|---|---|
| Model composition | | |
| • Non-hydrogen atoms | 8848 | 8848 |
| • Protein residues | 1212 | 1212 |
| • Ligands | 4 (GTP), 4 (MG) | 4 (GTP) |
| Resolution estimates (Å) | | |
| • d model | 3.8 | 4.0 |
| • d FSC model (0/0.143/0.5) | 3.1/3.3/6.9 | 3.8/3.9/7.8 |
| Validation | | |
| • MolProbity score | 1.89 | 1.90 |
| • Clashscore | 12.23 | 14.86 |
| Rotamer outliers (%) | 0.45 | 0.56 |
| Ramachandran plot | | |
| • Favored (%) | 95.76 | 96.59 |
| • Allowed (%) | 4.24 | 3.41 |
| • Disallowed (%) | 0 | 0 |

**Figure S4**

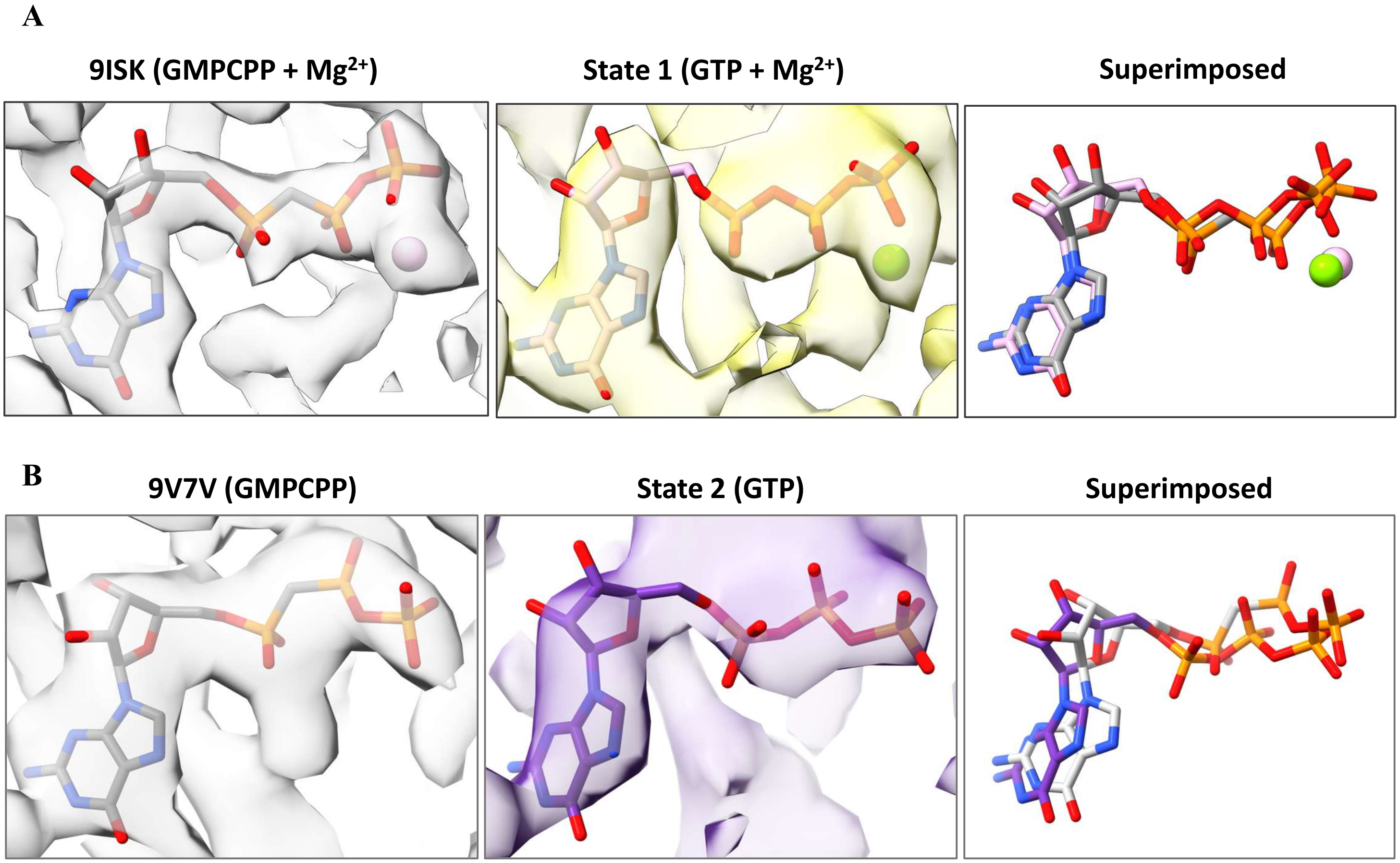


**Figure S4.** A. Comparison of GTP orientation between SmFtsZ state 1 (current study) and earlier published cryo-EM structures of *K. pneumoniae* FtsZ PDB 9ISK and EMD-60837. B. Comparison of GTP orientation between SmFtsZ state 2 (current study) and earlier published cryo-EM structures of Odinarchaeota FtsZ PDB 9V7V and EMD-64825

**Figure S5**

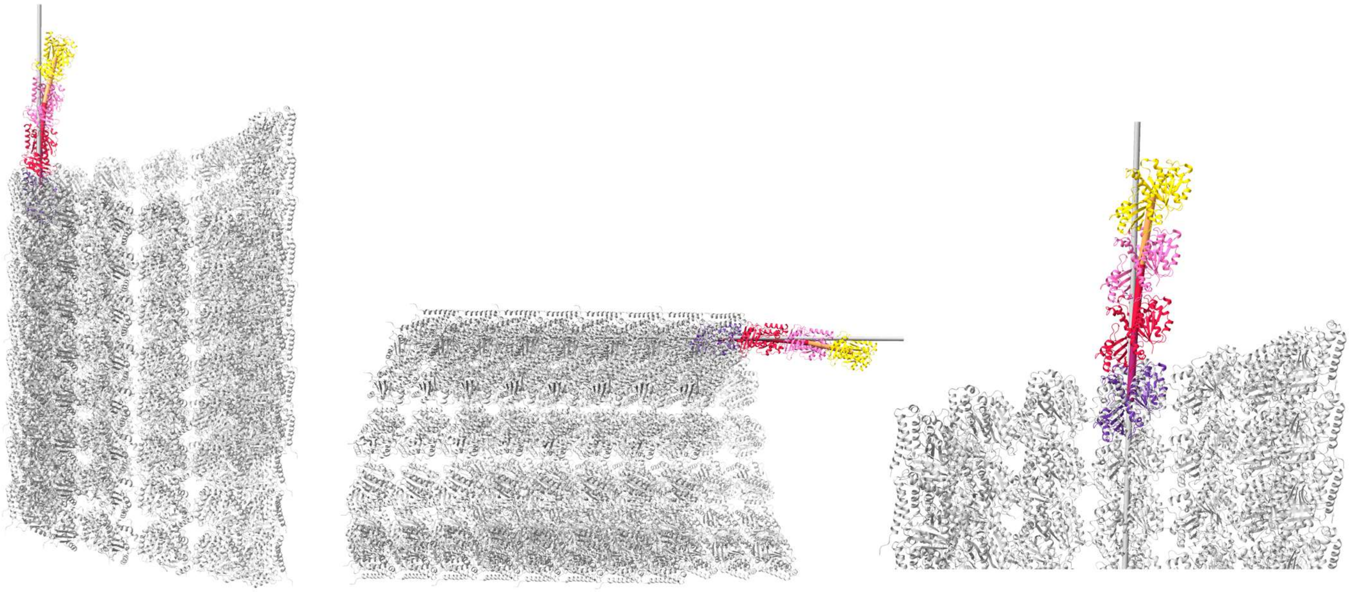

**Figure S5. Comparison of curvature of FtsZ protofilament with respect to the microtubule lumen vs exterior.** The superimposed filaments are shown from three viewing angles.

Figure S6

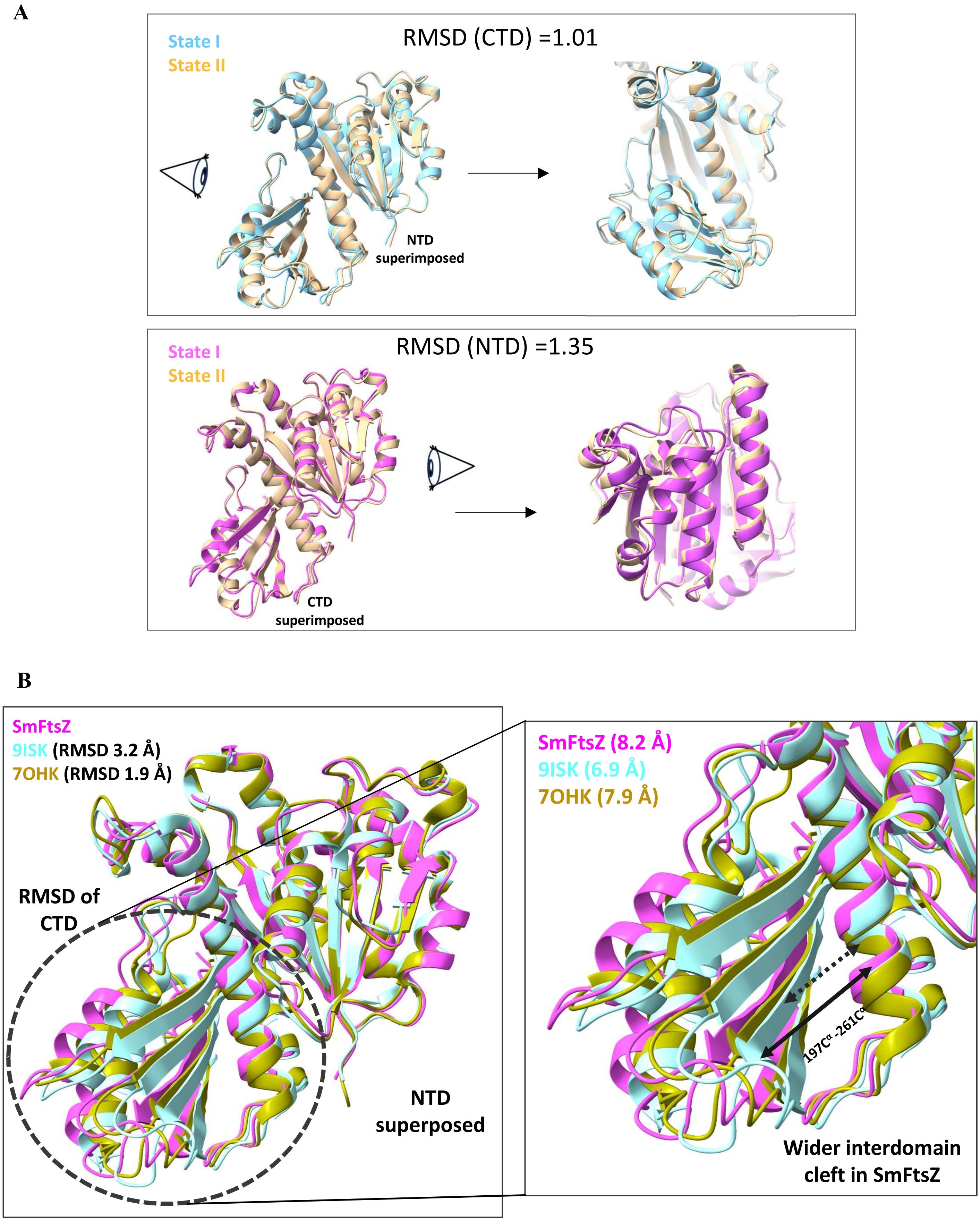


**Figure S6. Comparison of structural flexibility of two structural states (state 1 and state 2) of SmFtsZ filament.** (A) RMSD calculation of CTD (top) and NTD (bottom) by superimposition of NTD and CTD respectively. B. (Left) Superimposition NTDs of SmFtsZ state 1 with existing high resolution cryo-EM structure (PDB 9ISK) and crystal structure (PDB 7OHK). CTD is marked with a dotted circle (Right) Magnified region from dotted area showing distances between H7 helix (197C$^{\alpha}$ residue) and CTD (261C$^{\alpha}$ residue).

**Figure S7**

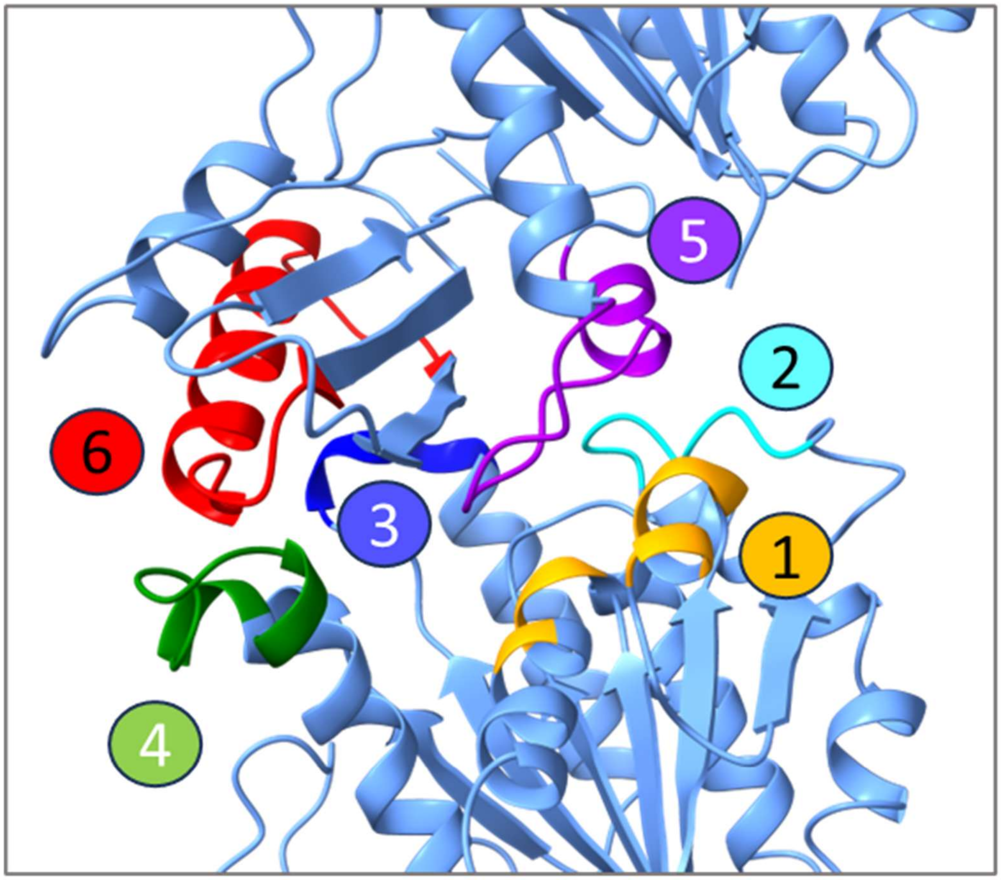


**Figure S7. Assignment of various interfacing zones and interactions at the longitudinal interface of the SmFtsZ.** Six different zones are shown in different colors: zone 1, yellow; zone 2, cyan; zone 3, blue; zone 4, green; zone 5, purple; and zone 6, red.

**Figure S8**

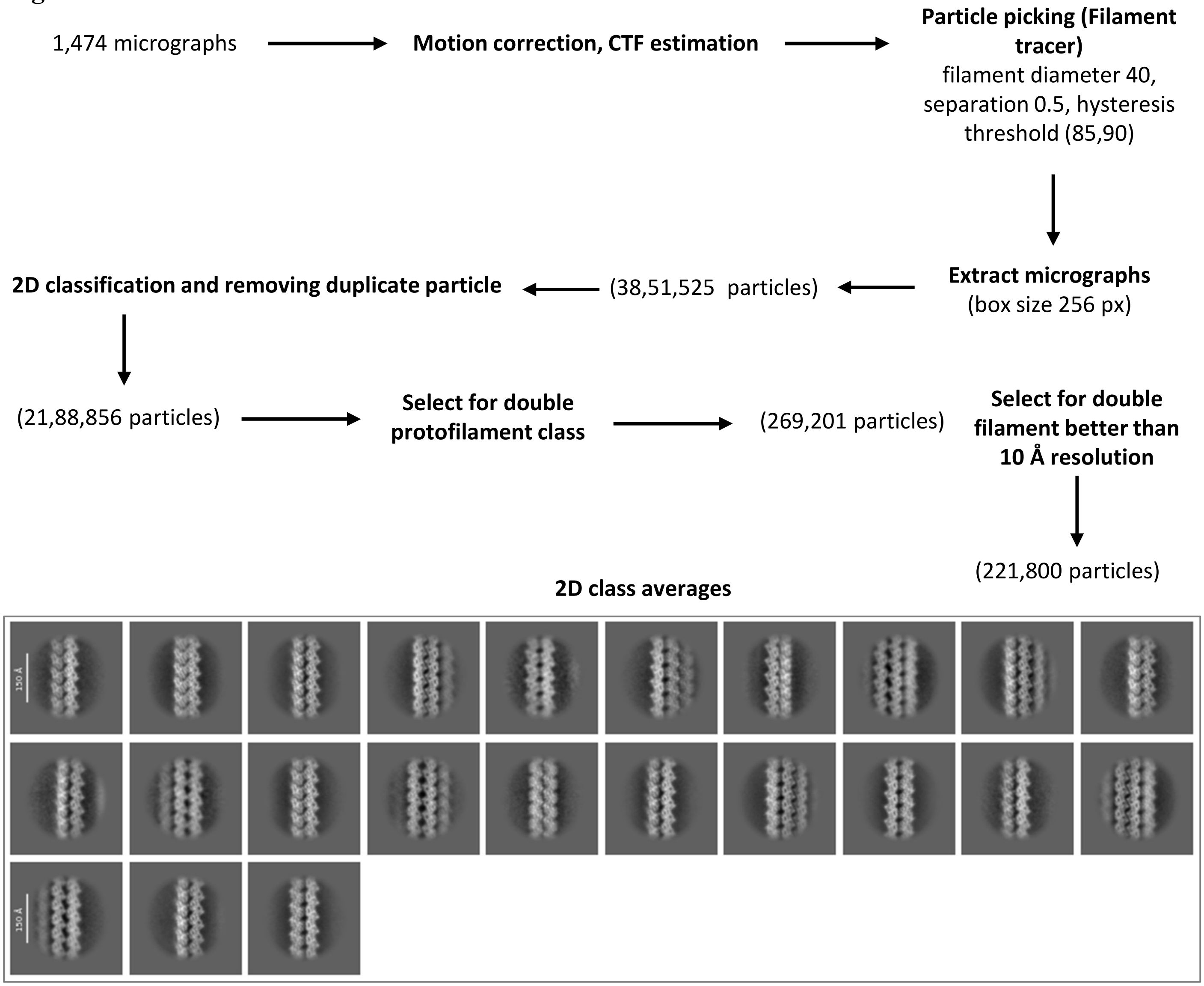


**Figure S8. 2D classification workflow of SmFtsZ double protofilament assembly**

**Figure S9**

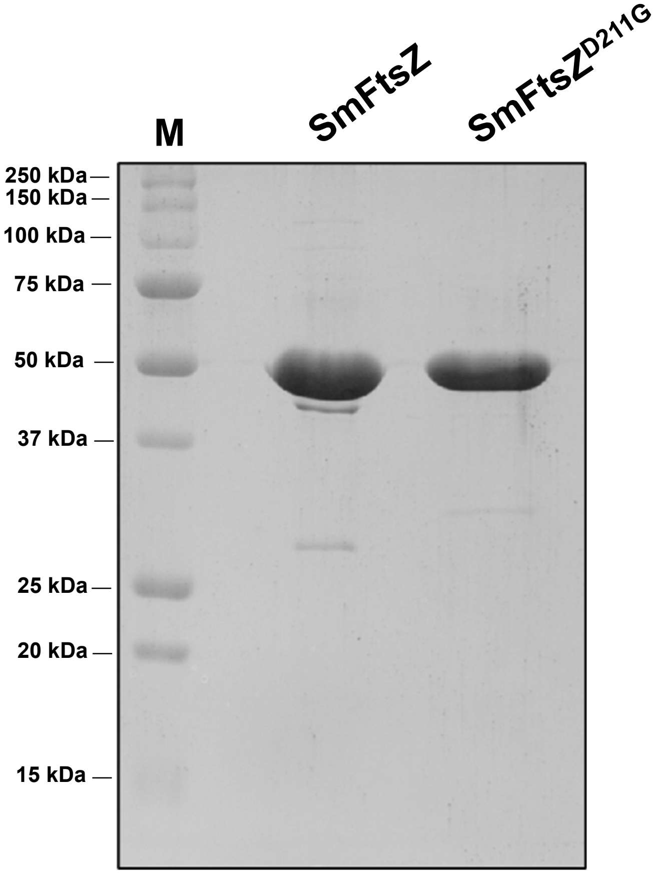


**Figure S9. Protein purity check gel**
Final purified and concentrated proteins were run on a 12% SDS PAGE for checking purity. First lane (M) corresponds to protein marker.

**Supplementary movie 1. 3D variability analysis of SmFtsZ curved state 1**

**Supplementary movie 2. 3D variability analysis of SmFtsZ curved state 2**

**Supplementary movie 3. Comparison of curvature between SmFtsZ and microtubule protofilament in a helical tube structure.**

**Supplementary movie 4. Dynamic and static zones are shown in SmFtsZ filament as a morph of state 1, state 2 and hypothetical straight model filament structures.**
